# Natural variation in tetrapyrrole biosynthetic enzymes and their regulation modify the chlorophyll mutant *Oy1-N1989*

**DOI:** 10.1101/2024.09.23.614405

**Authors:** Amanpreet Kaur, Rajdeep S. Khangura, Brian P. Dilkes

## Abstract

Tetrapyrroles are macrocyclic compounds present in and required for all life on Earth. Mutants with defects in tetrapyrrole pathway enzymes can be used to uncover natural variation in this pathway and study pathway regulation. We report here the effects of the *Oy1-N1989/+* mutation, a semi-dominant allele of subunit I in the Mg-chelatase enzyme with reduced chlorophyll biosynthesis, on global gene expression and morphological traits. Coordinate regulation of the tetrapyrrole pathway was observed as transcriptional feedback regulation of genes in the tetrapyrrole pathway in *Oy1-N1898*/+ mutants. Natural variation in the wild-type allele at *oy1* modulated the severity of the impact of *Oy1-N1989/+* on gene expression. We also studied the effects of previously identified cis-acting expression variation at *oy1* in wild-type plants. The seventy transcripts encoding biosynthetic enzymes in the tetrapyrrole pathway exhibited similar transcriptional co-regulation in response to cis-variation in *oy1* transcript accumulation in wild-type plants as observed in the RNAseq of *Oy1-N1989*/+ mutants. This demonstrated that the coordinate regulation of the pathway also occurs during physiologically-relevant variation in OY1 abundance. Cis variants at seven tetrapyrrole pathway genes were linked to variation in chlorophyll accumulation in *Oy1-N1989/+* mutant or wild-type plants. Analysis of trans-acting transcriptional variation by eGWAS detected multiple transcriptional hotspots, which disproportionately affected the expression of a subset of tetrapyrrole pathway genes, indicating that some genes are repeated targets of transcriptional regulation. The trans-regulatory hotspots coordinately regulated this pathway and may work to limit the accumulation of phototoxic intermediates.

**Author Summary:** The mutant allele and interactions with the genetic background determine the severity of mutant phenotypes. We used *Oy1-N1989/+* mutants, affecting Mg-chelatase subunit I, to identify natural variants in the tetrapyrrole pathway affecting transcriptional regulation and variation in mutant traits. By integrating GWAS and transcriptomics, we identified similar feedback regulation of porphyrin and chlorophyll metabolism in mutants as well as in wild-type plants affected by natural variation. This is the first demonstration of natural variation affecting transcriptional feedback in the tetrapyrrole pathway under normal conditions. Using GWAS as a regulatory discovery tool, we identified unexplored regulatory mechanisms in a pathway that is critical for energy capture from the sun and chemical energy utilization in all living things and multiple human diseases.

## Introduction

Tetrapyrroles are macrocyclic compounds, pigments of life, involved in various biological processes fundamental to all life on earth [1,2]. The diverse roles of tetrapyrroles are defined by the oxidation state of the macrocycle, the composition of sidechains, and their ability to chelate metals, including Magnesium, Iron, Cobalt, or Nickel. Tetrapyrroles in plants include chlorophyll, heme, siroheme, and phytochromobilin. All these compounds are synthesized from glutamyl tRNA in a multi-branched pathway (Figure 1).

**Figure 1.**
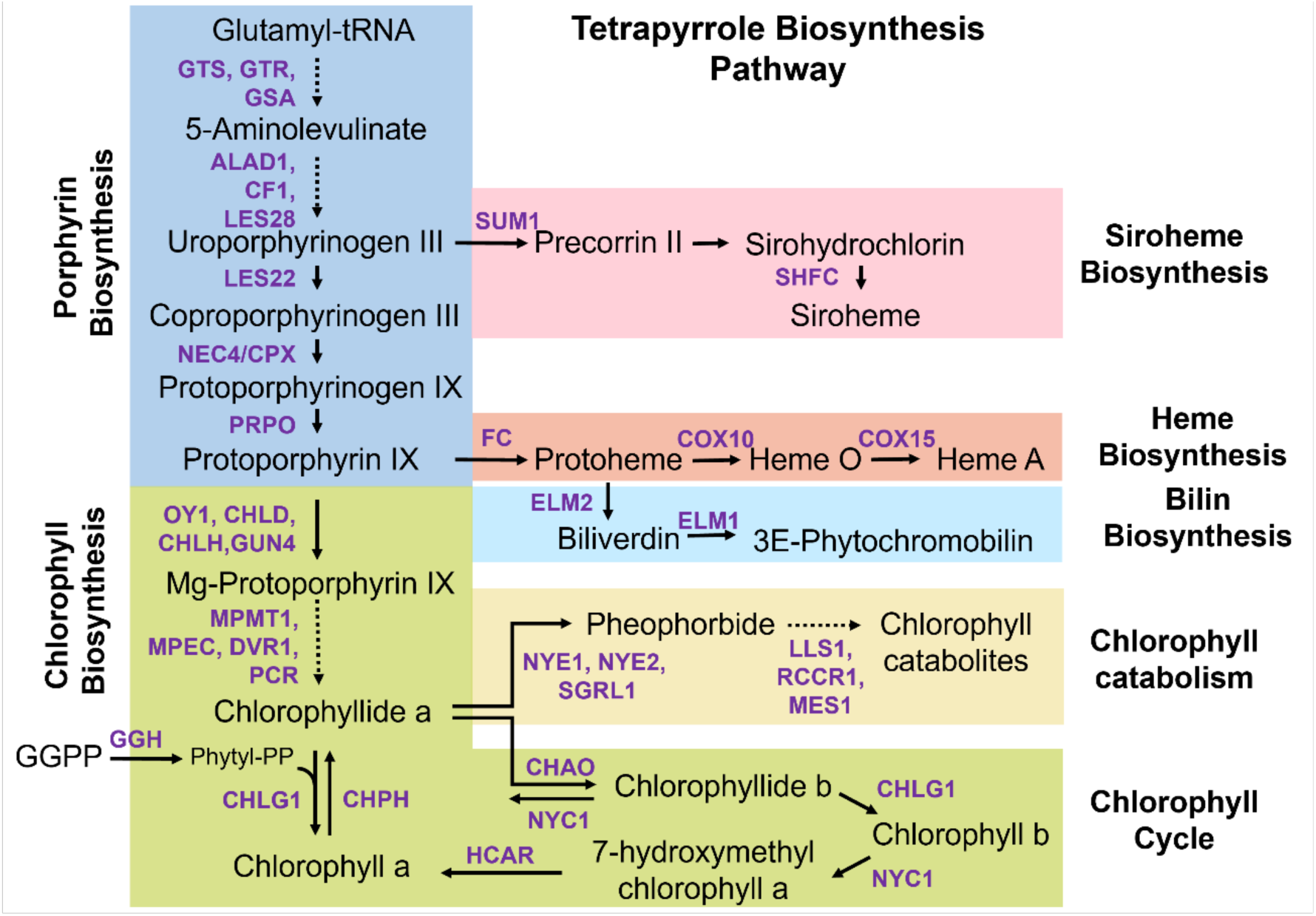
Schematic of tetrapyrrole biosynthetic pathway in maize. The enzymes catalyzing the steps in the pathway are depicted in purple. The details of abbreviations are in Supplemental Table S5.

Balancing the demands between branches of this pathway requires complex interplay and regulation. One aspect of this regulation is coordinate transcriptional repression or activation of tetrapyrrole biosynthetic genes. Light signaling, circadian rhythm, phytohormones, and environmental (biotic and abiotic) stresses mediate transcriptional regulation in the tetrapyrrole biosynthetic pathway [3,4]. Several transcription factors that coordinately affect the expression of genes encoding steps in tetrapyrrole metabolism are described in Arabidopsis [3,5]. ALA biosynthesis marks the first committed step in the biosynthesis of tetrapyrroles. The conversion of glutamyl tRNA to 5-aminolevulinate (ALA) is catalyzed by three enzymes: glutamyl-tRNA synthetase (GTS), glutamyl-tRNA reductase (GTR) and glutamate-1-semialdehyde 2,1-aminomutase (GSA). Studies of transcriptional regulation of the pathway have focused on glutamyl tRNA reductase (GTR), encoded by HEMA1 in Arabidopsis, as it controls the biosynthesis of ALA that serves as the precursor to all the branches of the pathway[3]. GTR activity is also regulated post-translationally through feedback inhibition by heme and interaction with a regulatory protein FLUORESCENT (FLU), which mediates the repression of ALA synthesis under fluctuating light conditions [5,6]. ALA is converted to uroporphyrinogen III by the action of three enzymes: ALA dehydratase (ALAD), porphobilinogen deaminase, and uroporphyrinogen-III synthase. These three steps are conserved in all living organisms, produce phototoxic intermediates, and are common to the biosynthesis of all tetrapyrroles [7]. Little is known about their regulation in plants. The tetrapyrrole pathway bifurcates at uroporphyrinogen-III into either siroheme biosynthesis or continues to protoporphyrin IX. Protoporphyrin IX is a common precursor for the chlorophyll and heme/bilin branches. The insertion of Fe^2+^ by Ferrochelatase (FC) is the first committed step for heme biosynthesis, and the insertion of Mg^2+^ to protoporphyrin IX by multimeric Mg-chelatase is the first committed step for chlorophyll biosynthesis. Mg-chelatase consists of three subunits, CHLI, CHLD, and CHLH, and requires the hydrolysis of ATP [8,9]. The activities of Mg-chelatase and Ferrochelatase are responsive to light signaling and concentrations of ATP. Mg-chelatase activity requires ATP, whereas Ferrochelatase is inhibited by ATP [10]. Additional regulation by protein complex subunits includes GUN4, which stimulates the activity of Mg-chelatase [11] and YCF54, which positively regulates Mg-protoporphyrin IX monomethylester cyclase [12–14]. Genes encoding steps in the chlorophyll branch, including the *chlh* gene encoding subunit H of Mg-chelatase, *gun4, chl27* encoding Mg-protoporphyrin IX monomethyl ester cyclase (MPEC), and *por* genes (*pora, porb, porc*) encoding protochlorophyllide oxidoreductases (PCRs) are known targets of transcriptional regulation [3,5]. The bilin pathway synthesizes phytochrome chromophore Phytochromobilin, a known regulator of the tetrapyrrole pathway [15,16]. Heme is metabolized by heme oxygenase, encoded by the maize *elm2* gene [17], to synthesize Phytochromobilin [17] (Figure 1).

Tetrapyrroles absorb photons and can convert light energy into chemical energy to the benefit and detriment of living things. While chlorophylls and bilin contribute to photosynthesis and energy production, some intermediates produced during the synthesis and breakdown of tetrapyrroles are photoreactive. These phototoxic intermediates can be excited by light to produce free radicals and reactive oxygen species that damage cells [18]. To avoid ROS accumulation, biosynthesis of the phototoxic intermediates in the tetrapyrrole pathway must be tightly controlled.

Mutants with defects at various points in the pathway can help us understand the functioning of the tetrapyrrole pathway and its regulation. Several mutants defective in tetrapyrrole metabolism have been identified and characterized in maize. Weak *Mutator*-induced maize *camouflage1* (*cf1*) mutants have decreased porphobilinogen deaminase activity and develop yellow non-clonal sectors on the leaves grown in diurnal light cycles but not on leaves grown in continuous light [19]. A loss of function allele in the same gene encodes the *necrotic3 (nec3)* mutant which has necrotic bands on leaves [20]). Semi-dominant alleles of *lesion22* (*les22*) encoded by one of the uroporphyrinogen decarboxylase paralogs of maize cause the light-dependent formation of necrotic lesions on the leaves due to accumulation of phototoxic uroporphyrinogen III [21]. A defect in coproporphyrinogen III oxidase in a recessive allele of *necrotic4*, *nec-t* leads to necrotic spots and yellow-green leaves [22]. These mutants have reduced chlorophyll contents and reduced accumulation of key intermediates of chlorophyll biosynthesis, including protoporphyrin IX, Mg protoporphyrin IX, and protochlorophyllide. Another mutant*, pale-green leaf (pgl),* containing missense mutation in a gene encoding magnesium-protoporphyrin IX monomethyl ester cyclase (MPEC) led to a chlorophyll-deficient phenotype in the leaves throughout the lifespan of the plant [23]. Multiple semi-dominant *Oil yellow1 (oy1)* mutants encode dominant-negative alleles of the subunit I of the Mg-chelatase enzyme and exhibit pale green leaf phenotype with low chlorophyll accumulation when heterozygous [24].

We previously demonstrated that combining natural and induced genetic variants can identify previously unknown regulation in this pathway. The semi-dominant allele, *Oy1-N1989,* encoded by a missense mutation in the Mg-chelatase subunit I, is a dominant-negative protein that poisons the Mg-chelatase complex. Using this mutant, we identified cryptic natural variation in the chlorophyll biosynthetic pathway that affected variation in mutant phenotypic severity [21–24]. Cis-regulatory natural variation at the *oy1* locus modified the chlorophyll contents of *Oy1-N1989/+* mutants [25]. In line crosses, the phenotypic consequences of *Oy1-N1989*/+ heterozygotes were enhanced by the *oy1*^Mo17^ allele, which accumulated less OY1 transcript, and suppressed by the *oy1*^B73^ allele, which accumulated more OY1 transcript. The severity of the effects of *oy1*^Mo17^ was evident in the greater reduction in chlorophyll content, lower photosynthetic rate, decreased carbohydrate accumulation, reduced stalk width, and delayed reproductive maturity as compared to mutants with *oy1*^B73^ allele [25,26]. In a GWAS study, natural cis-regulatory expression variation in the accumulation of OY1 transcript also encoded a strong modifier of *Oy1-N1989/+* mutant severity [25]. The connection between expression GWAS (eGWAS) and phenotypic trait variation and the ability of the natural variants at *oy1* to modify the phenotype of the *Oy1-N1989* mutant led us to explore the gene expression consequences of this mutant and natural variation in the entire tetrapyrrole pathway.

In this study, we explored transcriptional regulation of the tetrapyrrole pathway in maize using the *Oy1-N1989* mutant and its modifiers. Differential gene expression analysis in mutants displayed coordinate regulation of the tetrapyrrole pathway and compensatory effects at multiple genes encoding enzymes in tetrapyrrole biosynthesis. We performed a pathway-level exploration via expression level GWAS. Natural variation at *oy1* acted in trans on the accumulation of transcripts encoding multiple tetrapyrrole biosynthetic enzymes in wild-type plants. This demonstrates that even without discernable morphological effects in wild-type plants, regulatory variants at the natural alleles are consequential and trigger detectable transcriptional feedback regulation. The eGWAS also detected many transcriptional hotspots. Decomposition of those hotspots to single SNPs allowed us to demonstrate that natural alleles coordinately regulate the tetrapyrrole pathway, some genes in the tetrapyrrole pathway are frequent targets for transcription regulation, and development alters the pattern of transcript co-regulation. These patterns suggest an explanation for the banding patterns visible on the *cf1* mutant of maize encoded by weak alleles of porphobilinogen synthase.

## Results

### Gene expression consequences of *Oy1-N1989* in two genetic backgrounds are correlated with their phenotypic severity

We have previously reported the morphological and developmental consequences of the enhancement and suppression of the *Oy1-N1989*/+ phenotype by the *vey1* QTL [25–27]. The *vey1* QTL likely encodes a cis-regulatory expression polymorphism at the *oy1* locus, a widespread variant in the maize germplasm [25]. The wildtype *oy1* allele derived from B73 (*oy1*^B73^) accumulates higher OY1 transcripts than the *oy1* allele derived from Mo17 (*oy1*^Mo17^). As a result, the *Oy1-N1989/oy1*^B73^ mutants accumulate less chlorophyll than their wild-type siblings, and the introgression of the *oy1*^Mo17^ allele in B73 background to create isogenic *Oy1-N1989/oy1*^Mo17^ mutants results in a further reduction in chlorophyll [25].

In the current study, we first explored the global transcriptional consequences of *oy1*^B73^ and *oy1*^Mo17^ alleles on *Oy1-N1989* mutants in isogenic B73 genetic background. For this experiment, a near-isogenic line b094 that carries homozygous *oy1*^Mo17^ introgression in B73 background and B73 itself were crossed with *Oy1-N1989/oy1*^B73^ pollen-parent to create F1 progenies [21] that segregated ∼1:1 for mutant and wild-type siblings (Figure 2A). The cross of b094 with *Oy1-N1989/oy1*^B73^ pollen-parent produced severely chlorotic *Oy1-N1989/oy1*^Mo17^:B73/b094 F1 mutants, hereafter called as *Oy1-N1989/oy1*^Mo17^:b094, and normal *oy1*^B73^*/oy1*^Mo17^:B73/b094 F1 wild-type siblings, hereafter called as *oy1*^B73^*/oy1*^Mo17^:b094 (Figure 2A). The cross of B73 with *Oy1-N1989/oy1*^B73^ pollen-parent produced suppressed chlorotic *Oy1-N1989/oy1*^B73^:B73 mutants and normal *oy1*^B73^*/oy1*^B73^:B73 wild-type siblings. The RNA-sequencing analyses were carried out on pooled triplicates of all four genotypes. Principal component analysis (PCA) of the global transcriptional changes showed that the wild-type samples from B73 and B73/b094 backgrounds were transcriptionally similar. In contrast, the suppressed *Oy1-N1989/oy1*^B73^:B73 and severe *Oy1-N1989/oy1*^Mo17^:b094 mutant samples formed two distinct groups (Figure 2B). The first principal component, PC1, captured this grouping and mirrored the loss of chlorophyll. The wild-type samples overlap for PC1, a small increased value was observed for the suppressed *Oy1-N1989/oy1*^B73^:B73 mutant samples, and a greater value was calculated from all replicates of the severe *Oy1-N1989/oy1*^Mo17^:b094 mutants.

**Figure 2.**
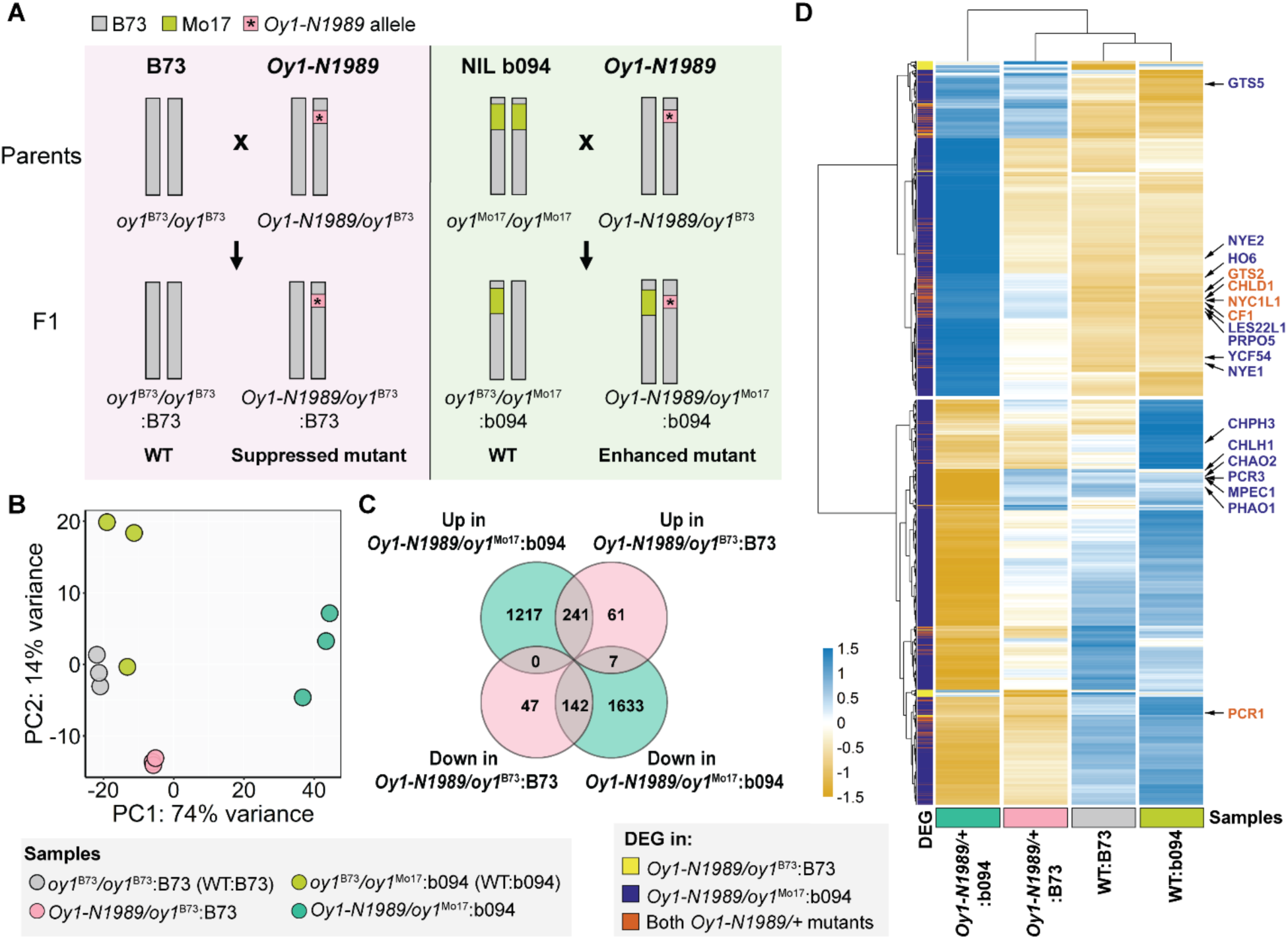
Transcriptomic analysis of *Oy1-N1989/+* mutants. A) The crossing scheme to generate isogenic F1 hybrids segregating 1:1 for wild-type plants and *Oy1-N1989/+* mutants in the B73 and B73 x b094 backgrounds; B) Principal component analysis (PCA) of transcriptomic data; C) Venn diagram showing number of genes up-regulated and down-regulated in suppressed *Oy1-N1989/oy1*^B73^:B73 and enhanced *Oy1-N1989/oy1*^Mo17^:b094 mutants as compared to their respective congenic wild-type controls; D) Heatmap showing normalized counts of transcripts differentially accumulated in either of the two *Oy1-N1989/+* mutants. The transcripts highlighted in red are differentially accumulated in both *Oy1-N1989/+* mutants and those in dark blue are only differentially accumulated in enhanced *Oy1-N1989/oy1*^Mo17^:b094 mutants.

Differential expression analysis identified 498 DEGs in *Oy1-N1989/oy1*^B73^:B73 compared to wild-type B73 siblings (Figure 2C). Of these 498 DEGs, transcripts of 309 were increased, and 189 were decreased in their abundance in the mutant. The number of DEGs was much higher in the enhanced *Oy1-N1989/oy1*^Mo17^:b094 mutants compared to their congenic wild-type siblings, with a total of 3240 DEG. Among these 3240 transcripts, 1458 were increased, and 1782 were decreased in accumulation in the enhanced mutants. In a comparison of the enhanced and suppressed *Oy1-N1989/+* mutants, 383 (77%) of the DEGs in the *Oy1-N1989/oy1*^B73^:B73 were also differentially expressed and altered in the same direction when the phenotype was enhanced by the *oy1^Mo17^* allele in *Oy1-N1989/oy1*^Mo17^:b094 (Figure 2C). Only seven (1.4%) of the DEGs common between *Oy1-N1989/oy1*^B73^:B73 and *Oy1-N1989/oy1*^Mo17^:b094 were expressed in opposite directions, demonstrating that the gene expression changes affected in the phenotypically milder background were similarly affected when the phenotype was enhanced.

The genes that were differentially expressed only in one background were also examined. We identified 2850 genes that were differentially expressed only in the enhanced *Oy1-N1989/oy1*^Mo17^:b094 mutant as compared to its wild-type siblings and 108 genes that were DEG only in the comparison between the suppressed *Oy1-N1989/oy1*^B73^:B73 mutant and its wild-type siblings. We explored the effect of enhancement of *Oy1-N1989/+* by *oy1*^Mo17^ by quantifying the effects on transcript levels for all DEGs in both the mutants compared to their respective wild-type siblings (Figure 2D). A total of 82% of the genes that comprised the DEG in either *Oy1-N1989/+* mutant comparisons were altered in the same direction in both the enhanced and suppressed mutants but did not pass statistical thresholds to be included as DEG. This provides molecular evidence that the enhanced impact of *oy1*^Mo17^ on *Oy1-N1989/+* has a quantitative effect on gene expression, much like the changes observed in chlorophyll content and plant growth of these mutants.

We investigated this further by comparing the magnitudes of expression effects between suppressed *Oy1-N1989/oy1*^B73^:B73 and enhanced *Oy1-N1989/oy1*^Mo17^:b094 mutants. To perform an aggregate analysis across global transcriptional dysregulation, a Z-score was calculated from normalized transcript counts for each gene in the DEG list. The Z-score provided a measure of the relative expression of the gene across our two mutants and wild-type samples. The similarity between expression regulation was quantified by averaging the Z-scores of all genes in the DEG list grouped by their expression direction, a value we refer to as an index [28,29]. Such parametric comparison allows the inference of the magnitude of the effect of each mutant condition on a gene expression pattern. The *Oy1-N1989/oy1*^B73^:B73 induced indices were calculated for up-regulated genes and repressed indices for the down-regulated genes in the *Oy1-N1989/oy1*^B73^:B73 mutant. The index for the genes induced by *Oy1-N1989/oy1*^B73^:B73 had a higher value in the enhanced *Oy1-N1989/oy1*^Mo17^:b094 mutants (Figure 3A). This pattern did not change when using the index values obtained with the genes induced by *Oy1-N1989/oy1*^Mo17^:b094 as well as with the genes repressed in *Oy1-N1989/oy1*^Mo17^:b094. The aggregate expression of genes measured as indices demonstrate that *Oy1-N1989/oy1*^Mo17^:b094 has a greater impact on expression level than *Oy1-N1989/oy1*^B73^:B73 (Figure 3B). To ensure robustness and control bias in our gene set selection, we calculated indices only using the genes that were significantly induced or repressed in both mutants. These “gold standard” DEG indices exhibit the same pattern, with greater aggregate values for the magnitude of gene expression changes in *Oy1-N1989/oy1*^Mo17^:b094 than in *Oy1-N1989/oy1*^B73^:B73 (Figure 3C). Thus, the findings of expression magnitude comport with the overall DEG numbers (Figure 2) and previous demonstration of its severe chlorotic phenotype [25]. This global transcriptional regulation data demonstrated that *Oy1-N1989/oy1*^Mo17^:b094 is an enhanced version of the *Oy1-N1989/oy1*^B73^:B73 phenotype and is not the result of a novel or synthetic effect of the induced and natural variation brought together in these materials.

**Figure 3.**
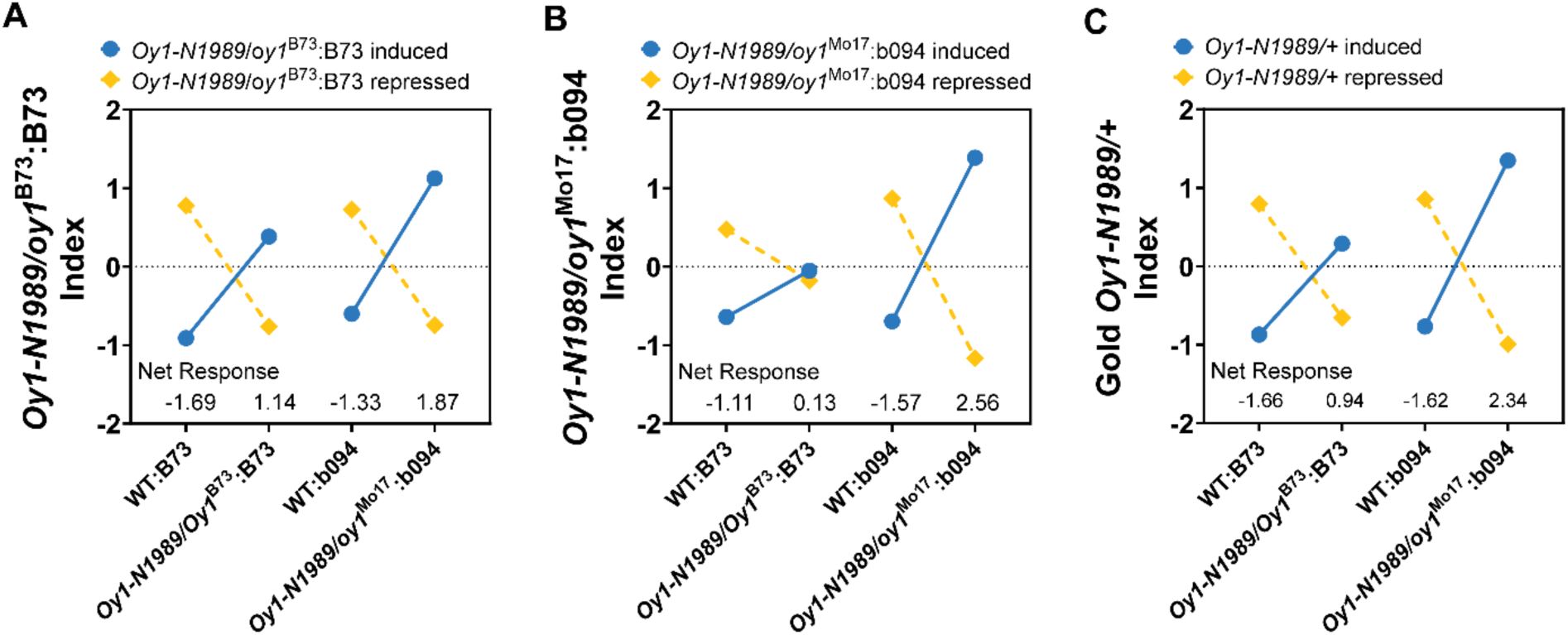
Comparison of expression effects in *Oy1-N1989/+* mutants and their wild-type controls using A) *Oy1-N1989/oy1*^B73^:B73 induced and repressed indices; B) *Oy1-N1989/oy1*^Mo17^:b094 induced and repressed indices; C) Gold *Oy1-N1989/+* induced and repressed indices. Gold indices represent genes that were differentially expressed in both *Oy1-N1989/+* mutants. The induced indices in each comparison are depicted by solid blue circle and repressed indices are depicted by a gold diamond. The index values provide relative comparison of expression effects across the four samples.

### Loss of Mg-chelatase activity in *Oy1-N1989* affects porphyrin and chlorophyll metabolism

The *Oy1-N1989/+* mutants carry a semi-dominant and dominant-negative allele of the *oy1* gene. This gene encodes the CHLI subunit of Mg-chelatase, the first committed step in chlorophyll biosynthesis that catalyzes the incorporation of Mg^2+^ into Protoporphyrin IX [24]. In our RNA seq experiments, OY1 (Zm00001d023536) was not differentially accumulated in *Oy1-N1989/+* mutants. The *Oy1-N1989/+* mutant backgrounds significantly increased the accumulation of transcripts encoding Mg-chelatase subunit D (CHLD1, Zm00001d013013) (Figure 2D). Transcripts encoding CHLH1 (Zm00001d026603) were significantly reduced in *Oy1-N1989/oy1*^Mo17^:b094 but not differentially accumulated in *Oy1-N1989/oy1*^B73^:B73. Transcripts of other genes encoding steps in chlorophyll biosynthesis and breakdown were also affected. Transcripts of *protochlorophyllide reductase1 (pcr1;* Zm00001d001820) were decreased in both the mutants. Transcripts of several other chlorophyll biosynthetic genes, including *Mg-protoporphyrin ester cyclase 1* (*mpec1*, Zm00001d008230), *pcr3* (Zm00001d013937), and homologs of chlorophyllide a oxygenase, CHAO2 and PHAO1 (Zm00001d011819 and Zm00001d042026), were decreased in the enhanced mutant. Thus, except for the CHLD1 transcript, DEGs within the chlorophyll biosynthetic pathway were reduced in *Oy1-N1989*/+ mutants. In addition, the transcripts of genes in the porphyrin pathway upstream of Protoporphyrin IX showed an increased accumulation in *Oy1-N1989/+* mutants (Figure 2D). Transcripts of *glutamate tRNA synthetase* (*gts2*, Zm00001d015037) and *camouflage1* (*cf1*, Zm00001d015366), which encodes a hydroxymethylbilane synthase, were increased in both the suppressed as well as enhanced mutants. In addition, enhanced mutants also accumulated transcripts of *glutamyl tRNA synthetase5* (*gts5*, Zm00001d048372), *lesions22-like1* (*les22l1*, Zm00001d011386) which encodes a uroporphyrinogen decarboxylase, and *protoporphyrinogen IX oxidase5* (*prpo5*, Zm00001d030962). Thus, defects in Mg-chelatase subunit I in *Oy1-N1989/+* mutants caused a decrease in expression of chlorophyll biosynthetic genes and increased expression of upstream porphyrin biosynthetic genes.

Gene expression patterns were less consistent at transcripts encoding steps in other branches of the tetrapyrrole pathway. Transcripts of the chlorophyll degrading enzyme CHLOROPHYLLASE3 (CHPH3/CHAO1, Zm00001d031934) were decreased in *Oy1-N1989/oy1*^Mo17^:b094 (Figure 2D). However, transcripts of other genes involved in chlorophyll catabolism, including *nyc1l1* (Zm00001d013651), *nye1* (Zm00001d021288), and *nye2* (Zm00001d006211) were DEG with increased expression in *Oy1-N1989/oy1*^Mo17^:b094 as compared to the wild type. Among the genes involved in chlorophyll degradation, only *nyc1l1* was a DEG in *Oy1-N1989/oy1*^B73^:B73 and showed increased accumulation. The genes in other branches of the tetrapyrrole pathway, including the siroheme, heme, and bilin biosynthesis, were not differentially expressed in either suppressed or enhanced *Oy1-N1989/*+ mutants (Supplemental Table S1).

To obtain a comprehensive view of the effect of *Oy1-N1989* on tetrapyrrole biosynthesis, we examined the expression differences at the maize homologs of all genes involved in porphyrin, chlorophyll, siroheme, heme, and bilin biosynthesis and chlorophyll degradation (Figure 4, Supplemental Table S5). For this purpose, a p-value ≤ 0.05 was used to identify genes with differences in gene expression, and the direction of the effect on expression was assessed. Of the 18 expressed genes encoding steps in porphyrin biosynthesis upstream of chlorophyll biosynthesis, the expression differences for 12 had p-value ≤ 0.05 in *Oy1-N1989/+* mutants in both the enhancing and the suppressing backgrounds. Both the mutants showed a higher accumulation of 11 out of these 12 differentially expressed transcripts (Figure 4, Supplemental Table S5). In addition to this set of shared transcripts, transcripts of *lesion22* (*les22*, Zm00001d029074) and *protoporphyrinogen IX oxidase2* (*prpo2*, Zm00001d003214) were differentially accumulated in the enhanced mutant. These results demonstrate the existence of a transcriptional feedback mechanism that responds to a mutation affecting Mg-chelatase activity to trigger the accumulation of transcripts of upstream porphyrin biosynthetic enzymes.

**Figure 4.**
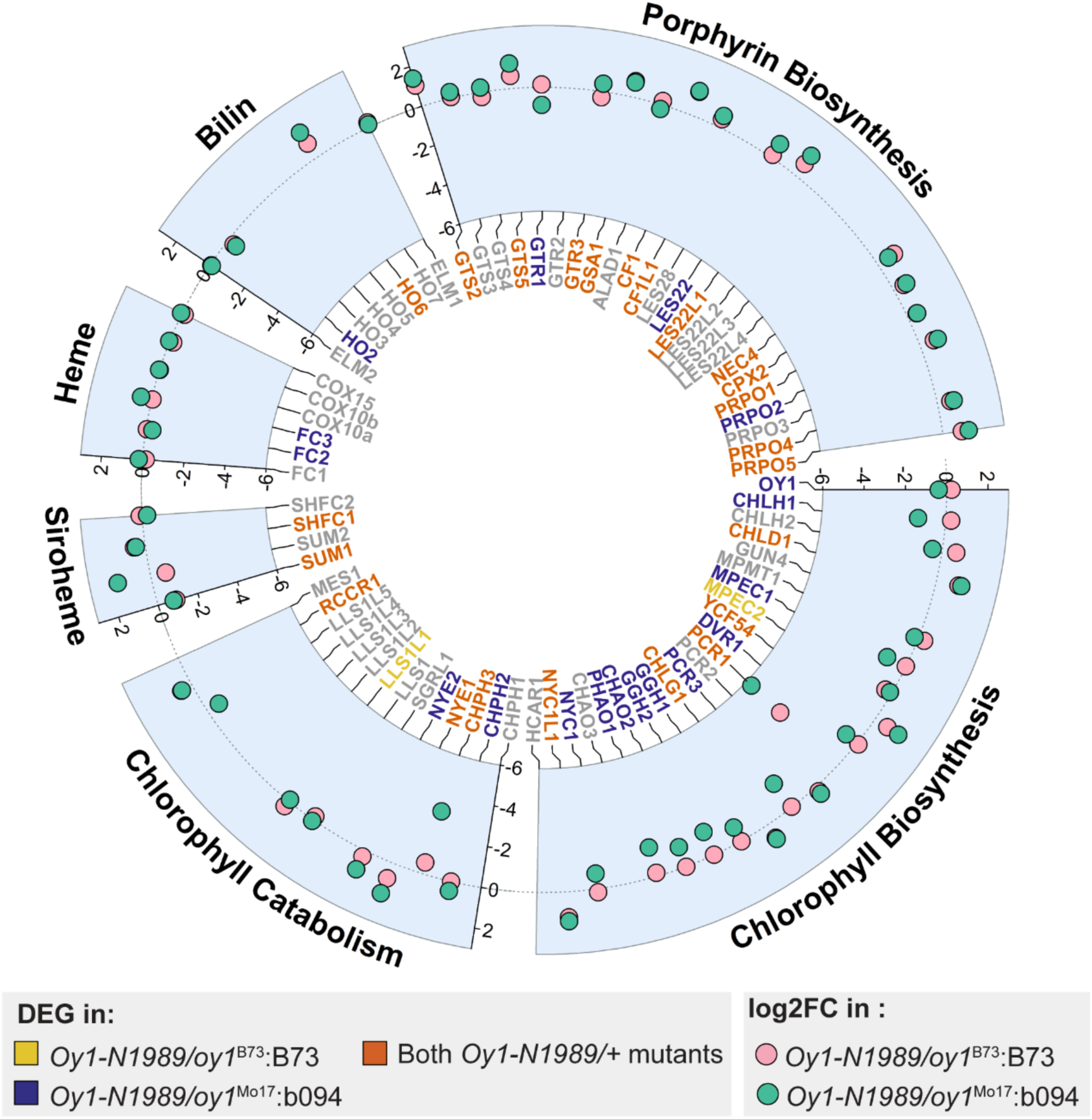
Effect of *Oy1-N1989* on expression of genes encoding steps in tetrapyrrole biosynthetic pathway. Circle plot shows log2 fold change of transcripts of genes encoding steps in tetrapyrrole biosynthetic pathway in *Oy1-N1989/oy1*^B73^:B73 (pink circles) and *Oy1-N1989/oy1*^Mo17^:b094 (green circles) mutants as compared to their respective wild-type controls. The transcripts highlighted in red were differentially expressed in both *Oy1-N1989/+* mutants at unadjusted p-value (p ≤ 0.05). Transcripts highlighted in yellow and dark blue were only differentially expressed in *Oy1-N1989/oy1*^B73^:B73 and *Oy1-N1989/oy1*^Mo17^:b094 respectively. Transcripts in grey were either not differentially expressed at unadjusted p-value cutoff of 0.05 or were not detected in our RNA-seq data.

A similar analysis was carried out on the 19 expressed genes encoding steps in the chlorophyll biosynthesis branch of the pathway. The transcripts of 15 out of 19 expressed genes in the chlorophyll biosynthesis pathway were differentially expressed at p-value ≤ 0.05 in *Oy1-N1989/oy1*^Mo17^:b094, and only six were differentially expressed in *Oy1-N1989/oy1*^B73^:B73. The enhanced mutants showed a reduced accumulation of 11 of 15 differentially expressed transcripts for chlorophyll biosynthetic genes. Five of these 11 transcripts were also repressed in the suppressed mutant background, but four were increased in their accumulation (Figure 4, supplemental Table S5). This demonstrates a case of epistasis where the allele at *oy1* alters the effect direction of transcript accumulation in the *Oy1-N1989* background. What mechanisms and transcription factors mediate this feedback is unknown, let alone how they could result in such a non-linear effect. A similar non-linear or opposing effect of the *oy1* allele has been previously observed for plant height, where a suppressing *oy1*^B73^ allele results in taller mutant plants compared to isogenic wild-type siblings [26]. In contrast, a severe *oy1*^Mo17^ allele results in shorter mutant plants than isogenic wild-type siblings. That said, it appears that severe loss of Mg-chelatase activity triggers reduced expression of chlorophyll biosynthetic enzymes and thus may contribute to the severe loss of chlorophyll in *Oy1-N1989/oy1*^Mo17^:b094 mutants.

Along with these coordinate changes in transcripts encoding tetrapyrrole pathway steps, transcripts of transcriptional regulators of chloroplast development and the tetrapyrrole pathway were differentially expressed in *Oy1-N1989/oy1*^Mo17^:b094 mutants. Transcripts of *golden plant2* (*g2*, Zm00001d039260) and *phytochrome-interacting factor4* (*pif4*, Zm00001d013130) were decreased in *Oy1-N1989/oy1*^Mo17^:b094 as compared to its wild-type siblings. Transcripts of *long hypocotyl5* (*hy5*, Zm00001d015743), *phytochrome1* (Zm00001d033799), and *phytochrome2* (Zm00001d013402) were increased in accumulation in the enhanced mutant. The negative regulators of GA-induced gene expression DELLA-transcription factors (D8, Zm00001d033680 and D9, Zm00001d013465) were decreased in accumulation in *Oy1-N1989/oy1*^Mo17^:b094. An analysis of GO enrichment for both RNA seq experiments is described in the supplementary materials (Supplemental Text and Supplemental Figure S1).

### Cis-acting regulatory polymorphisms at tetrapyrrole pathway genes impact chlorophyll content in *Oy1-N1989/+* mutants

The severity of the mutant phenotypes in *Oy1-N1989/+* F1 hybrids is influenced by natural variation in the B73 and Mo17 alleles at the wild-type copy of *oy1*. To explore this further, we performed an eGWAS for the expression of *oy1* in three leaf tissues (L3Base, LMAD, LMAN) in maize diversity panel obtained from a previous study [30]. We defined cis polymorphisms as SNPs detected within 50 kb of the gene. Cis-acting polymorphisms affecting expression variation at *oy1* were detected in mature leaf tissues harvested during the day (LMAD) and night (LMAN) (Figure 5A) but not in the L3Base data set. We identified 26 cis SNPs in LMAD and 10 in LMAN associated with variation in OY1 transcript accumulation at a p-value of less than 10^-4^ (Supplemental Table S6). We evaluated these 36 cis SNPs for their impact on chlorophyll accumulation in *Oy1-N1989/+* mutants using data from our previous study, where we crossed the maize association panel with the *Oy1-N1989/oy1*^B73^ mutants [25]. Consistent with our previous studies [25,31], cis polymorphisms at *oy1* were significantly associated with chlorophyll accumulation in mutants and chlorophyll content ratios and differences for each F1 family (Figure 5B, Supplemental Table S7). The alleles associated with greater accumulation of OY1 were consistently associated with increased chlorophyll content in the mutant (Supplemental Table S7). No significant associations were identified for OY1 transcript accumulation and wild type chlorophyll contents. Thus, this exposed cryptic phenotypically impactful variation as the *Oy1-N1989* allele was required for these phenotypic effects, as expected due to previously described epistasis [26].

**Figure 5.**
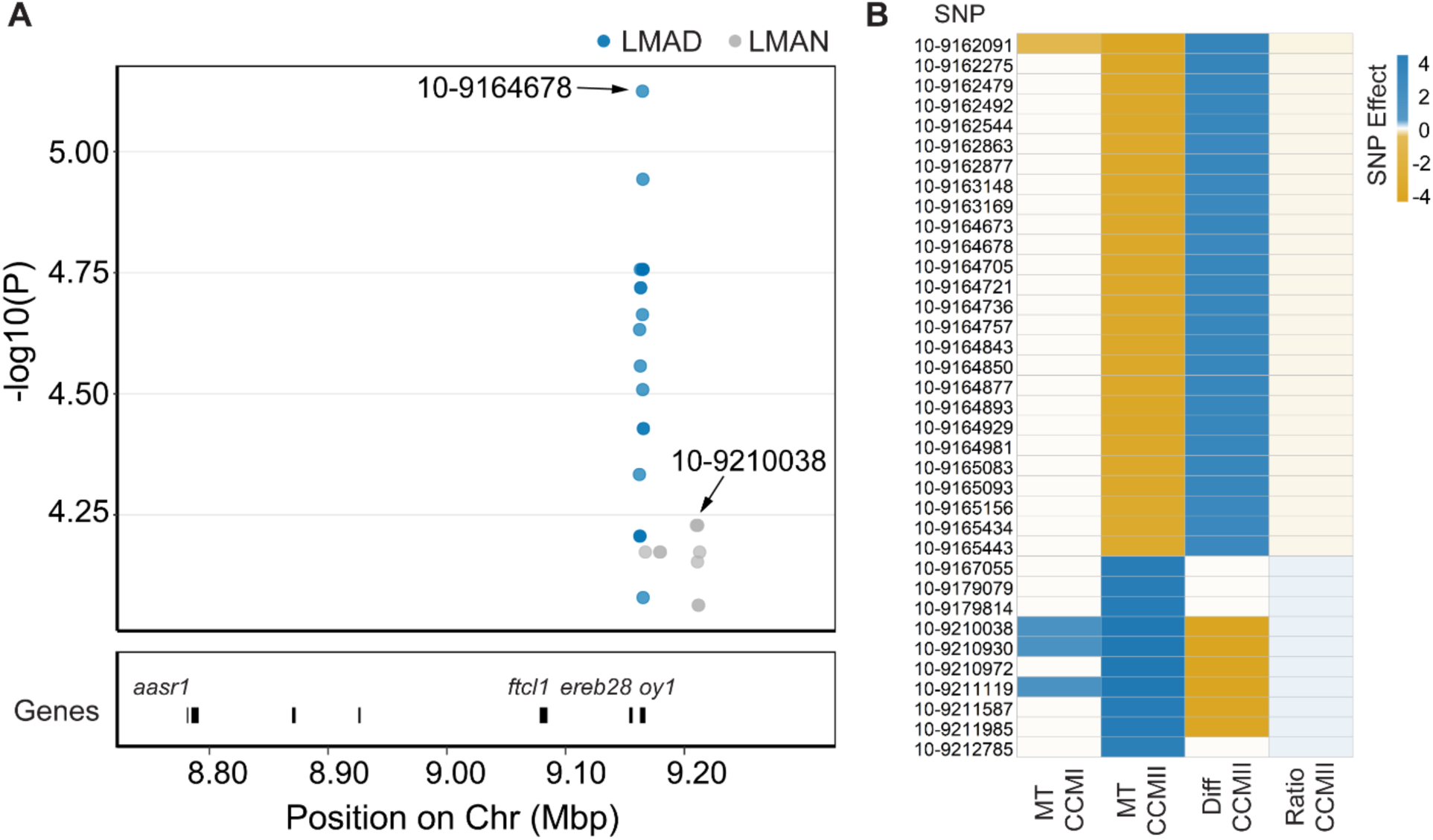
Cis-acting polymorphisms at *oy1* impact chlorophyll content in *Oy1-N1989/+* mutants. A) Cis polymorphisms affecting the expression of *oy1* in mature leaf tissues, LMAD (Blue dots) and LMAN (grey dots) at p-value < 1e-4; B) Effect of cis SNPs on chlorophyll accumulation traits. Blue color indicates a positive effect of SNP on phenotypic trait and gold color indicates a negative effect on the trait value at p-value < 1e-4. White color indicates that association between the SNP and the phenotypic trait was not significant at p-value < 1e-4.

To determine if variation in the expression of other tetrapyrrole pathway genes could affect variation in chlorophyll content, we performed a similar analysis using all genes encoding steps in tetrapyrrole biosynthesis. Across three leaf tissues (L3Base, LMAD, LMAN), 70 of 76 tetrapyrrole pathway genes were expressed in the maize association panel [30]. Expression GWAS was carried out for all genes, and all associations at p-value ≤ 1e-4 were retained for further analysis. At these criteria, 57 genes, including *oy1*, had cis polymorphisms affecting gene expression variation in at least one of the three tissues (Supplemental Figure S2, Table 1, and Supplemental Table S6). Like the analysis performed with cis SNPs at *oy1*, we intersected cis-acting SNPs affecting the accumulation of tetrapyrrole biosynthetic transcripts with the SNPs affecting variation in chlorophyll content. We obtained the greatest association overlap between chlorophyll content and transcript counts from mature leaves during the day (LMAD) (Supplemental Figure S3, Supplemental Table S7). None of these associations were as strong as the association with *oy1*. Nevertheless, cis-regulation at *chlh1*, which encodes subunit H of the Mg-chelatase complex, was associated with the mutant CCMI, mutant CCMII, CCM ratio, and CCM difference traits (Supplemental Figure S3, Supplemental Table S7). Increased accumulation of CHLH1 was associated with increased chlorophyll content in the mutant. Like *oy1*, the cis variation at *chlh1* did not significantly affect chlorophyll content in wild-type siblings. Similar relationships between transcript abundance and CCM traits were observed at two paralogs of *protochlorophyllide reductase* (*pcr1* and *pcr3*) that encode enzymes in chlorophyll biosynthesis (Supplemental Figure S3, Supplemental Table S7). SNPs affecting the reduced expression of *pcr1* in LMAD, LMAN, and L3Base also reduced mutant CCMII. SNPs affecting increased expression of *pcr3* in LMAD were associated with increased mutant CCMI and ratio CCMI. Among the genes in the porphyrin pathway, only cis polymorphisms at *les22*, encoding a uroporphyrinogen decarboxylase, were significantly associated with mutant chlorophyll content. The alleles associated with increased accumulation of LES22 transcripts in LMAD and LMAN showed a reduction of chlorophyll content in mutants. A higher transcriptional activity encoding a chlorophyll precursor like *les22* is expected to increase chlorophyll content and not lower it. This inverse relationship likely indicates a feedback regulation affected by cis variation at *les22*. Cis polymorphisms linked to reduced expression of *chlorophyllase2 (chph2)*, a chlorophyll degrading enzyme, were significantly associated with higher chlorophyll accumulation in both mutant and wild-type plants. This inverse relationship between chlorophyll catabolism and chlorophyll content is consistent with the expectation. Cis polymorphisms associated with a homolog of a heme biosynthetic gene, *ferrochelatase2* (*fc2*), were associated with mutant chlorophyll content (CCMI, CCMII) and chlorophyll contents of mutants normalized to wild-type siblings (Ratio CCMII and Difference CCMII) (Supplemental Figure S3, Supplemental Table S7). The alleles linked to an increase in the expression of *fc2* negatively impacted the chlorophyll content in mutants.

**Table 1.** Summary of cis variation in transcript abundance detected by association testing of SNPs within 50 kb of the seventy genes encoding steps in the tetrapyrrole biosynthetic pathway.

| Tissue | Genes with cis eQTLs | Cis SNPs at p-value < 1e-4 | Unique cis SNPs at p-value < 1e-4 |
| --- | --- | --- | --- |
| L3 Base | 46 | 13953 | 13767 |
| LMAD | 49 | 11392 | 11203 |
| LMAN | 52 | 14760 | 14538 |

### Trans-regulatory natural variation repeatedly targets the same tetrapyrrole metabolism genes and results in co-regulation

We further explored natural variation in the expression of the genes encoding the steps in the tetrapyrrole pathway to include trans-acting loci. SNP-transcript associations using the transcript abundance of all 70 genes in the tetrapyrrole pathway were identified through eGWAS (Table 3). This identified multiple trans-acting associations within 250kb of the *oy1* locus that affected multiple tetrapyrrole metabolism transcripts. We identified 62 SNPs linked to the *oy1* locus associated with the accumulation of 28 transcripts encoding steps in tetrapyrrole metabolism in the base of the third leaf (Supplemental Table S8). Ten of these 62 SNPs affected the transcript abundance of at least two genes, and in all cases, the trans-regulatory effect displayed the same effect direction for all affected genes (Figure 6, Table 2, Supplemental Table S8). Thus, variation at the *oy1* locus, which encodes subunit I of Mg-Chelatase, triggers transcriptional regulation that results in the coordinate regulation of genes encoding steps in the tetrapyrrole pathway. The eight genes encoding transcripts affected by these ten SNPs encoded steps in porphyrin and/or chlorophyll branches of the tetrapyrrole pathway (Figure 6). Transcript levels of five of these eight genes were increased (p-value < 0.05) in both *Oy1-N1989/+* mutants in our RNA-seq experiments (Figure 6). We identified 253 SNPs at the *oy1* locus linked in trans to the expression of 25 tetrapyrrole pathway genes in LMAD (Supplemental Table S8). Of the 253 SNPs, 63 affected at least two genes (Supplemental Figure S4). Similarly, 63 SNPs linked to the *oy1* locus were associated with expression variation for 17 tetrapyrrole metabolism genes in LMAN. However, each of these SNPs only affected single genes. Together with the DEG analysis (Figure 2, Figure 3, Figure 4), GWAS strongly suggests that a transcriptional regulatory mechanism(s) affects feedback regulation in response to the loss of chlorophyll and demonstrates that this regulation is also active in wild-type plants. In this regard, it is worth noting that these trans-acting SNPs linked to *oy1* are more than 50kb from the gene itself and may indicate examples of local trans effects [32]. Consistent with this interpretation, each of the SNPs in Table 2 affects the accumulation of OY1 transcript in the same direction as the other tetrapyrrole genes and not in the opposite direction as would be expected for homeostatic compensation as observed in our RNA-seq experiments (Figure 4, Supplemental Table S5).

**Figure 6.**
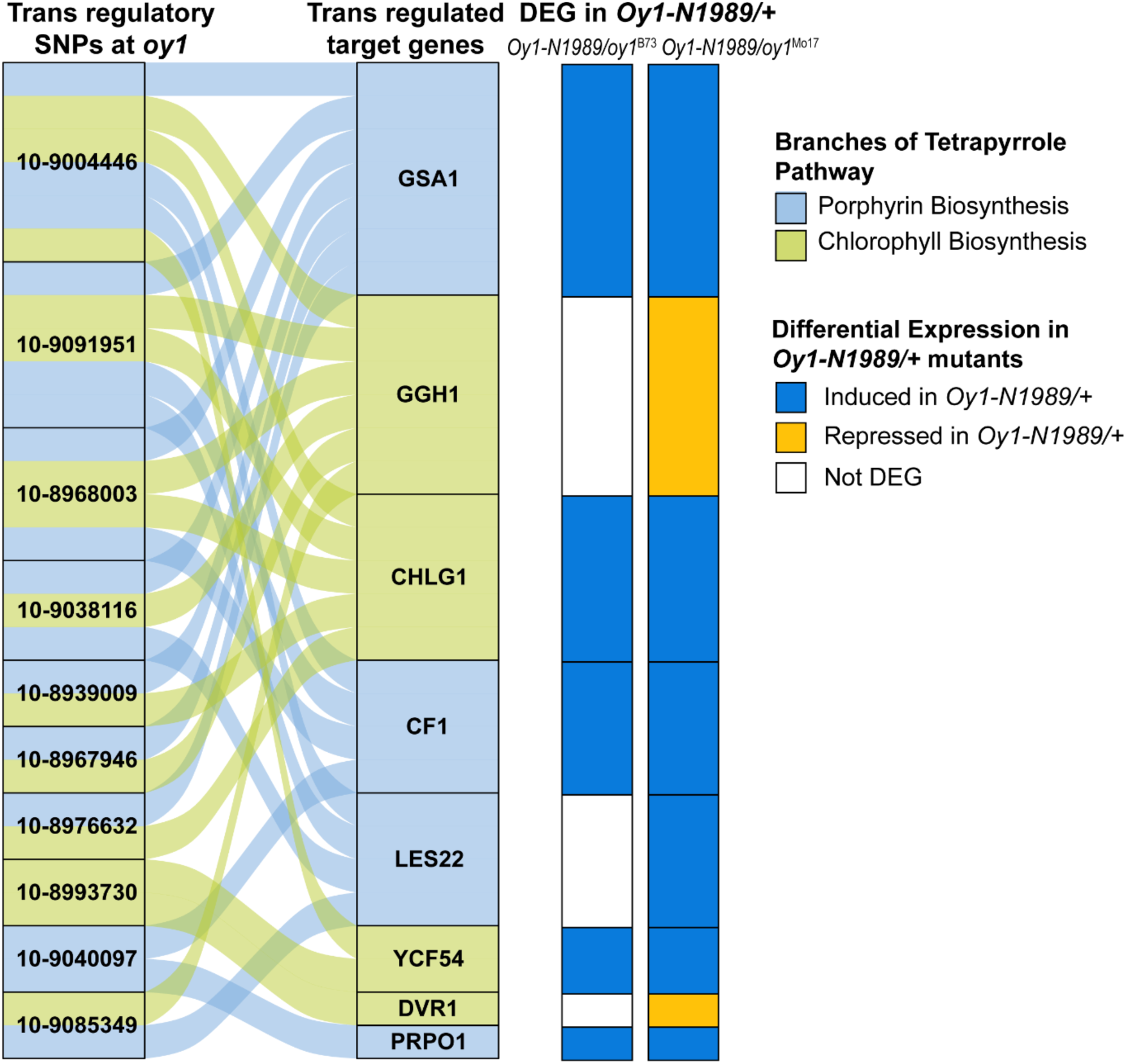
The *oy1* locus regulates the accumulation of transcripts encoded by porphyrin and chlorophyll biosynthetic genes. SNPs within a 250kb region around *oy1* associated in trans with transcripts of at least two genes encoding steps in tetrapyrrole pathway in base of third leaf. Light blue and green colors indicate the branches of the tetrapyrrole pathway. The panels on the right depict the direction of effect of *Oy1-N1989* on transcript accumulation in suppressed and enhanced *Oy1-N1989/+* mutants as compared to their respective congenic wild types at unadjusted p-value (p ≤ 0.05). Dark blue color indicates genes induced in the mutant and gold color indicates genes repressed in the mutant.

**Table 2.** Trans eQTLs at *oy1* locus linked to three or more genes in the tetrapyrrole biosynthesis pathway.

| Chr-Position<sup>a</sup> | Tissue<sup>b</sup> | Cis Effect on OY1 | eGWA OY1 p-value | Trans-regulated genes | Pathway<sup>c</sup> | Trans effect | Trans p-value |
| --- | --- | --- | --- | --- | --- | --- | --- |
| 10-8968003 | L3Base | -0.08 | 7.6E-05 | GSA1 | Porphyrin | -0.92 | 4.7E-05 |
|  |  |  |  | CF1 | Porphyrin | -0.49 | 6.3E-05 |
|  |  |  |  | GGH1 | Chlorophyll | -0.02 | 2.0E-05 |
|  |  |  |  | CHLG1 | Chlorophyll | -1.27 | 1.9E-05 |
| 10-9004446 | L3Base | -0.08 | 5.9E-05 | GSA1 | Porphyrin | -0.92 | 1.5E-05 |
|  |  |  |  | CF1 | Porphyrin | -0.48 | 3.2E-05 |
|  |  |  |  | LES22 | Porphyrin | -0.004 | 9.7E-05 |
|  |  |  |  | YCF54 | Chlorophyll | -0.20 | 2.5E-05 |
|  |  |  |  | GGH1 | Chlorophyll | -0.02 | 5.2E-05 |
|  |  |  |  | CHLG1 | Chlorophyll | -1.16 | 3.6E-05 |
| 10-9038116 | L3Base | -0.07 | 3.4E-04 | GSA1 | Porphyrin | -0.846 | 9.69E-05 |
|  |  |  |  | LES22 | Porphyrin | -0.004 | 4.68E-05 |
|  |  |  |  | GGH1 | Chlorophyll | -0.020 | 3.97E-05 |
| 10-9091951 | L3Base | -0.08 | 5.7E-04 | GSA1 | Porphyrin | -0.919 | 6.06E-05 |
|  |  |  |  | CF1 | Porphyrin | -0.488 | 6.99E-05 |
|  |  |  |  | LES22 | Porphyrin | -0.004 | 9.64E-05 |
|  |  |  |  | GGH1 | Chlorophyll | -0.022 | 4.15E-05 |
|  |  |  |  | CHLG1 | Chlorophyll | -1.231 | 4.37E-05 |
| 10-9020908 | LMAD | 0.70 | 2.1E-04 | GUN4 | Chlorophyll | 0.87 | 8.1E-05 |
|  |  |  |  | MPEC1 | Chlorophyll | 11.79 | 3.3E-05 |
|  |  |  |  | NYC1L1 | Chlorophyll | 0.19 | 5.8E-05 |

<sup>a</sup> Position in maize genome version 4; <sup>b</sup> tissue type defined in text <sup>c</sup> branch of the tetrapyrrole pathway see Figure 1

**Table 3.** Summary of trans-acting polymorphisms detected by expression GWAS of the seventy genes encoding steps in the tetrapyrrole biosynthesis pathway in three leaf tissues.

| Tissue | Trans-eGWAS<br>SNPs<br>below<br>p<1e-4 | Hot SNPs<sup>a</sup> | Top SNPs<sup>b</sup> | Hot Top<br>SNPs<sup>c</sup> | Trans<br>eQTL<br>hotspots<sup>d</sup> |
| --- | --- | --- | --- | --- | --- |
| L3Base | 385931 | 408 | 42917 | 55 | 338 |
| LMAD | 410722 | 33 | 47457 | 2 | 206 |
| LMAN | 519193 | 27 | 50862 | 3 | 220 |

<sup>a</sup> Hot SNPs affect trans-eGWAS at p-value <1e-4 for eight or more genes;<sup>b</sup> top SNPs were defined within exclusive windows of 250kb as the lowest p-value trans-acting SNP for each gene in each locus; <sup>c</sup> hot top SNPs encode the top SNP for eight or more genes; <sup>d</sup> hotspots were defined as 40kb windows encoding top SNPs that affected eight or more transcripts.

Using the eGWAS data for all 70 tetrapyrrole genes, we annotated trans-acting eQTL hotspots. We defined trans-regulatory hotspots in three ways. First, we annotated trans-acting eQTL at the SNP level, an analysis that permitted the detection of coordinate regulation of the tetrapyrrole pathway. We identified all trans-acting SNPs that affected the expression of eight or more tetrapyrrole pathway genes at a threshold of p-value < 10^-4^. We refer to these as “hot SNPs.” Though the threshold for each individual SNP is permissive, if we consider gene expression independent across all genes in each sample (which it is not, see discussion) exceeding a p-value < 10^-4^ threshold and affecting the expression of at least eight different genes from the set of 70 tetrapyrrole pathway genes should happen 9.8 x 10^-23^ times. We tested ∼2.4 x 10^7^ SNPs on this gene set, for which a Bonferroni threshold for alpha < 0.05 would be 2.1 x 10^-9^. By requiring expression effects at eight genes to qualify as a “hot SNP” this is far more stringent than a Bonferroni alpha < 0.05 and every “hot SNP” exceeds this threshold by 14 orders of magnitude. A second approach was taken that increases the stringency of this method further. We filtered the eGWAS results to retain only the top SNP (lowest p-value), affecting the expression of each gene within a window of 250kb. Single nucleotide positions that were the top SNP for eight or more genes were considered “hot top SNPs.” Third, we adopted a more traditional hotspot definition that used genomic windows to annotate regions with an unexpectedly large number of trans-acting eQTL as the criteria to define a hotspot locus. To identify loci affecting the expression of multiple genes, we used 40kb windows as our locus size. There are 62,500 40kb windows in the maize genome. To create a stringent definition of “hotspot”, we processed the GWAS output for each of the 70 pathway genes retaining only the top SNPs (lowest p-value) associated with gene expression in 250kb windows and used these as inputs for the hotspot calculation (Supplemental Table S9). To be a hotspot a 40kb window had to contain top SNPs associated with the expression of 8 different tetrapyrrole pathway genes. For the tissue with the greatest number of SNPs (LMAN), the average number of top SNPs per gene, rounding up, is 727 (50,862 Top SNPs across the 70 genes; Supplemental Table S9). At this rate, the proportion of windows containing a Top SNP for the average gene is 0.0116. The likelihood of an event with frequency of 0.0116 occurring at least 8 out of 70 genes is 1.63 x 10^-6^. Calculated in this way, 0.1 windows are expected to contain a hotspot calculated this and we observed 220 for LMAN, exceeding this number by three orders of magnitude. The other two tissues have fewer top SNPs (Table 3) and as a result we would expect even fewer windows to contain a hotspot, and yet observe 206 and 338 hotspots in LMAD an dL3 Base, respectively, exceeding this expectation by many orders of magnitude. To be more conservative and take into consideration the non-random distribution of SNP effects across genes due to the underlying biology, we calculated a False Discovery Rate by permutation. The number of associations per gene were tallied and assigned randomly to bins and the number of associations per bin were determined for 1000 permutated data sets. The average number of bins with eight or more associations was 0.137 per permutation, delivering a FDR for the observed set of 220 hotspots in LMAN as 6.2 x 10^-4^. False detection rates calculated by the same procedure for the 206 observed in LMAD was 3.6 x 10^-4^ and the 338 observed in L3 base was 1.6 x 10^-4^. Thus, all approaches presented here, whether tested as alleles at the SNP level or considered as loci and tested as genomic windows, detected far more trans-regulatory hotspots than expected by chance. This demonstrates pervasive heretofore unappreciated transcriptional co-regulation of the porphyrin pathway operating homeostatically within normal physiological ranges.

We identified 408 hot SNPs in the data from L3Base, 33 in LMAD, and 27 in LMAN (Table 3). Among these were 55 hot top SNPs in L3Base, two in LMAD, and three in LMAN. (Table 3, Supplemental Table S10). In L3Base, some transcripts were affected by more hot top SNPs than others demonstrating that some genes are more likely to be targets of trans-regulatory variation than others. For example, *cf1* had the greatest number of hot top SNPs affecting its accumulation (44 out of 55), indicating that the regulation of this gene was pervasive. A similar number (41 or 55) of hot top SNPs affected the expression level of *oy1*. As the hot top SNPs all affect multiple genes, we analyzed the allelic effect directions on all affected transcripts to explore pathway co-regulation by these natural variants (Figure 7).

**Figure 7.**
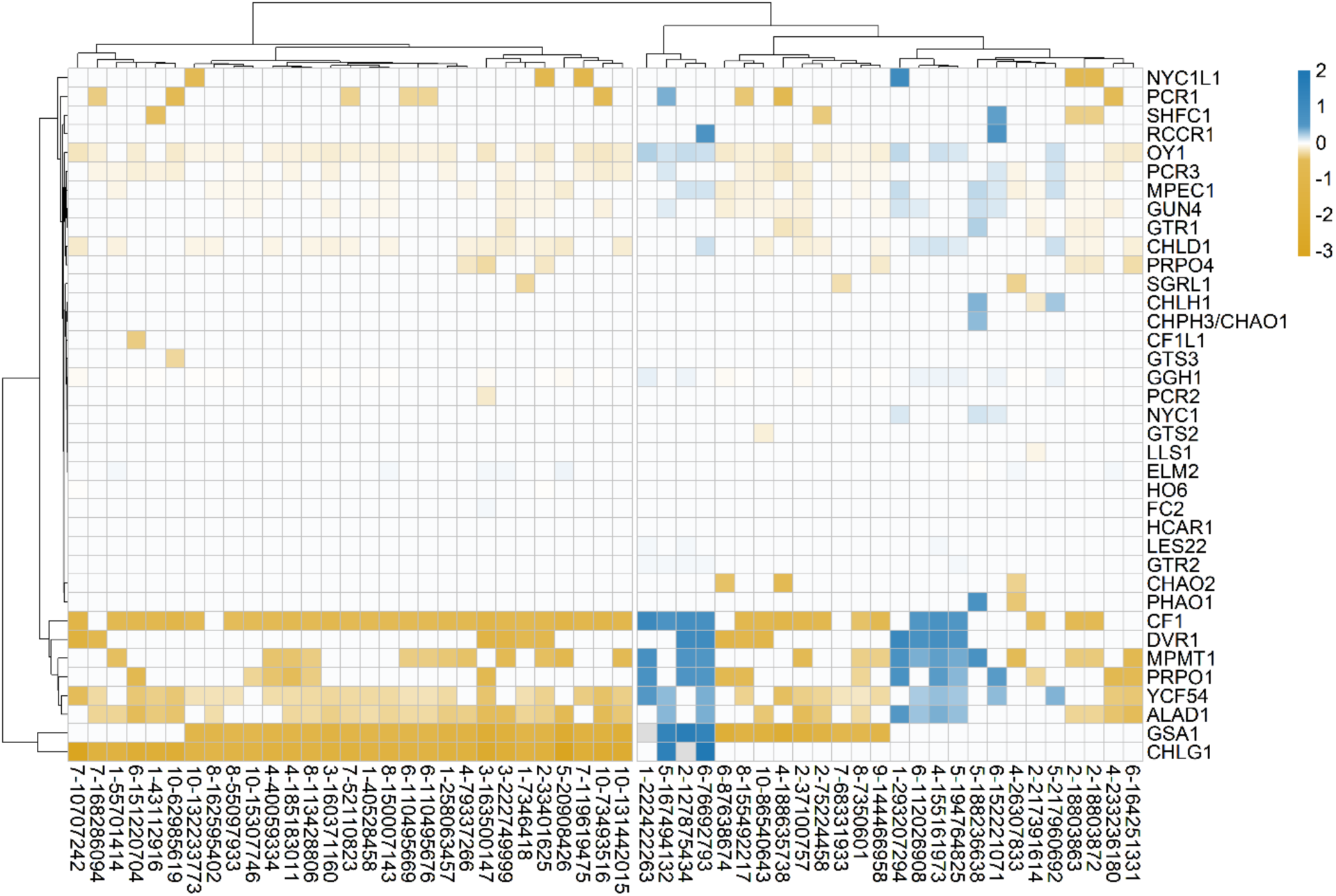
Effect of hot top SNPs on expression of associated genes in base of third leaf (L3Base). The color bar represents SNP effects. Blue color indicates a positive effect of SNP on gene expression and gold color indicates a negative effect on the expression at p-value < 1e-4. White indicates the effects were not significant at p-value <1e-4.

Remarkably, all 55 hot top SNPs in L3Base had consistent directional effects on SNP-transcript associations with core porphyrin and chlorophyll biosynthetic genes (Figure 7, Supplemental Table S10). For instance, SNP 8-150007143 in L3Base modified the expression of 14 tetrapyrrole pathway genes (Supplemental Table S10). The allele at this position affected the expression of 13 of these genes in the same direction, including *gtr2*, *gsa1*, *alad1*, *cf1*, *les22*, *chld1*, *oy1*, *gun4*, *mpec1, ycf54*, *pcr3*, *ggh1*, and *chlg1,* demonstrating that it alters a transcriptional regulatory pathway that coordinately regulates the chlorophyll and upstream porphyrin branches of this pathway. The only divergent effect direction at this hot top SNP (8-150007143) was with *elm2,* which encodes heme oxygenase involved in heme catabolism and the bilin branch of the pathway. This may identify a regulatory mechanism that permits coordination between the heme and chlorophyll pathways. For example, transcriptional regulation of porphyrin and chlorophyll biosynthesis may be coordinated through a change in the abundance of the phytochrome chromophore [33,34] or may result from heme oxygenase’s role in retrograde signaling [35].

In mature leaf collected at night (LMAN), the effect direction of all associations comprising the three hot top SNPs also resulted in coordinated directional effects on all transcripts encoding porphyrin and chlorophyll biosynthetic genes. In LMAD, however, allele effects were reversed at one of the two hot top SNPs for the transcript encoded by *cf1*. This hot top SNP, SNP 10-10925524, decreased the transcript accumulation of genes encoding steps in chlorophyll biosynthesis, including *oy1*, *chld1*, *gun4*, *mpmt1*, *mpec1*, *ycf54*, *dvr1*, *pcr3,* and the upstream porphyrin pathway gene *alad1*, but was associated with increased transcript levels of *cf1* (Supplemental Table S10). This, and the overwhelming occupancy of *cf1* effects in the hot top SNPs in L3Base, indicates a previously unknown regulation of this pathway at the porphobilinogen synthase step that may stop the accumulation of phototoxic intermediates, all of which occur after the step encoded by *cf1* while permitting continued operation of later steps.

Any 40kb genomic region that contained top SNPs affecting the expression of eight or more transcripts in the tetrapyrrole pathway was annotated as a trans-eQTL hotspot. This identified 338 trans-eQTL hotspots in L3Base, 206 in LMAD, and 220 in LMAN (Table 3, Supplemental Table S10). L3Base exhibited the greatest number of hotspots and a strong bias for trans-regulated genes (Figure 8), similar to that described using the hot top SNP approach (Figure 7 and Supplemental Table S10). Trans-eQTL hotspots affected genes encoding steps in all branches of the tetrapyrrole pathway. However, the genes encoding steps in the porphyrin and chlorophyll branches were the most common targets of hotspot variation (Figure 8). Similar results were observed in the mature leaf tissues (Supplemental Figure S5, Supplemental Figure S6). In addition, some tissue-specific effects on regulation were observed, such as the inclusion of chlorophyll catabolism genes among the repeated targets of trans-regulatory hotspots in mature leaf samples, which were consistent with the expression pattern of these genes.

**Figure 8.**
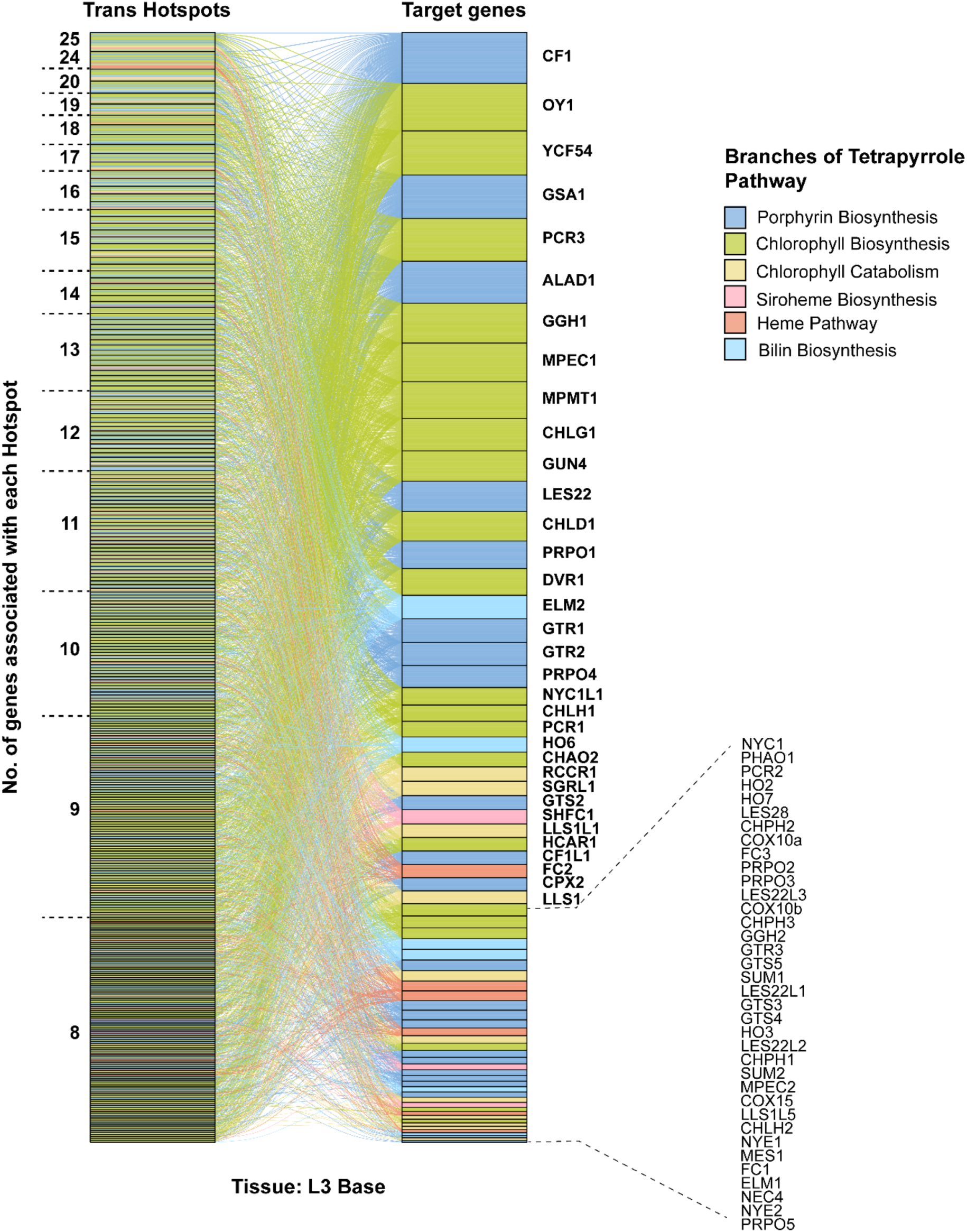
Trans-regulatory hotspots repeatedly target genes in porphyrin and chlorophyll branches of tetrapyrrole biosynthetic pathway. Trans eQTL hotspots associated with the transcripts of eight or more genes encoding steps in tetrapyrrole biosynthesis pathway in L3Base within a 40 kb window. The colors indicate different branches of tetrapyrrole biosynthetic pathway.

The trans-eQTL hotspots encoded a striking proportion of the overall trans-regulatory variation affecting the tetrapyrrole pathway. Seventeen genes had 10% or more of their trans-regulatory SNPs encoded within hotspots using the L3Base expression data (Supplemental Table S11). These included the transcripts from *gtr2*, *gsa1*, *alad1*, *cf1*, *les22*, *prpo1*, *oy1*, *chld1*, *gun4*, *mpmt1*, *mpec1*, *dvr1*, *ycf54*, *pcr3*, *ggh1*, *chlg1,* and *elm2.* Among the genes associated with trans-eQTL hotspots in LMAD, the transcripts of three genes, *gun4, mpmt1,* and *sgrl1,* had 10% of their trans-regulatory SNPs within hotspots (Supplemental Table S11). In LMAN samples, the transcripts encoded by *cf1* and *mpmt1* genes had more than 10% of their trans-regulatory SNPs encoded within hotspots (Supplemental Table S11). As the plants sampled for leaf three and mature leaves are different biological replicates, the finding that the same genes are repeatedly affected by trans-eQTL hotspots in different tissues strongly suggests that coordinated transcriptional regulation is a feature of these pathways.

We then tested whether the divergent regulation exhibited at CF1 compared to the rest of the pathway in mature leaves sampled during the day (Figure 7, Supplemental Table S10) was also a feature of these hotspots. Effect direction tests are possible with individual SNPs but not with windowed hotspot loci. To test this within the hotspots, we tabulated all SNPs within the hotspots that affected the CF1 transcript and at least two other genes. In LMAD, ten SNPs met these criteria, and without any exception, all SNPs affected CF1 transcript accumulation in the opposite direction of other porphyrin and chlorophyll pathway genes (Supplemental Table S10). In LMAN and L3Base, 15 and 116 SNPs affected the CF1 transcript and at least two different genes, respectively. In LMAN and L3Base, CF1 was coordinately regulated with other pathway genes at all SNPs, indicating that the time of day and the developmental stage of the leaf determined this phenomenon (Supplemental Table S10). This result demonstrates a change in the transcriptional feedback mechanisms or consequences in mature leaf tissue during the day relative to the two others. As *cf1* encodes the last step before the production of phototoxic intermediates [19,22,36] this may indicate transcriptional feedback to prevent the accumulation of these metabolites during the day.

### Chlorophyll accumulation is affected by a subset of trans-eQTL hotspots affecting the tetrapyrrole pathway

We compared the SNPs detected within these hotspots and the SNPs affecting variation in chlorophyll accumulation in the F1 association mapping experiment with *Oy1-N1989*. Again, we tabulated all overlaps that exceeded an arbitrary p-value ≤ 10^-4^ for each SNP-trait association. Given the number of hotspot SNPs (Table 3) and eight traits, we detected more associations than expected in all three expression sets (3 SNPs in L3Base, 4 in LMAD, and 7 LMAN), equivalent to a false detection rate of < 0.35 corresponding to 10 true detections of loci affecting both expression and phenotypic trait effects and five false overlaps for these QTL (Supplemental Table S12). An experiment-wide threshold of FDR ≤ 0.05 identified a single SNP from the expression hotspots in each tissue that affected both a hotspot and chlorophyll accumulation variation. In L3Base, this identified a hotspot resident SNP, 5-144452049, as associated with higher chlorophyll contents of the wild-type F1 plants and increased difference in CCM between wild-type and mutant siblings. This SNP was associated with a lower accumulation of the ELM2 transcript. We examined the effect of this SNP on chlorophyll content in the mutants. In our previous study, an *elm1* loss of function mutant synergistically enhanced the *Oy1-N1989*/+ phenotype [31]. Consistent with the synergistic interaction between *elm1* loss-of-function and *Oy1-N1989*/+, the allele at SNP 5-144452049 that decreased the abundance of ELM2 transcript decreased the CCM of *Oy1-N1989*/+ mutants (MT_CCMI: b=-0.71, p=0.019; MT_CCMII: b=-1.70, p-value=0.036). The opposite effect of 5-144452049 on the CCM measurements from mutant and wild-type siblings provides additional evidence that the heme/bilin and chlorophyll pathways engage in meaningful regulatory crosstalk affected by natural variation in maize. In LMAD, the hotspot resident SNP, 5-211181311, that passed the FDR threshold decreased the transcript abundance of LES22L2 and increased mutant CCMI and ratio CCMI traits. The role of the *les22*-like genes in maize is unknown. We explored the phylogenetics of these genes, as they were not previously investigated, and determined that they derive from duplication of uroporphyrinogen decarboxylase in the green plant lineage (Viridiplantae) before the divergence of the lineages leading to *Chlamydomonas* and the land plants and both lineages have retained members of both duplicates for more than a billion years (Supplemental Figure S7). In maize and Arabidopsis, one clade contains *les22* of maize and *heme2* of Arabidopsis, while the sister clade contains at least four paralogs in maize, *les22-like1* through *les22-like4*, and *heme1* of Arabidopsis. The vast majority of the tetrapyrrole pathway enzymes from Arabidopsis can complement their respective yeast mutant, but notably, neither HEME1 nor HEME2 of Arabidopsis can complement the yeast mutant defective in this step [37]. The SNP that passed the FDR < 0.05 threshold in the overlap with the LMAN hotspots was on chromosome 5 at 155197361 bp and coordinately affected the gene expression *gsa1*, *mpmt1*, and *dvr1* as well as the earlier of the two chlorophyll measures in the *Oy1-N1989/+* F1 association mapping data with effects on mutant CCMI, ratio CCMI, and difference CCMI. The allele associated with lower expression of these three genes reduced the chlorophyll content in mutant siblings (Supplemental Table S12), consistent with our hypothesis that this hotspot regulates chlorophyll accumulation via enzyme accumulation. These findings illuminate a gap in our understanding of the uroporphyrinogen decarboxylases of the green plant lineage, encoded in maize by *les22* and four ancient paralogs. They also highlight that while there are cases of significant overlaps between phenotypic impacts of the expression covariation in this pathway, the overwhelming majority of detected trans-regulatory hotspots neither affect the accumulation of chlorophyll in wild-type plants nor adjust the phenotype in response to the limitation on chlorophyll biosynthesis affected by the *Oy1-N1989* mutant.

### Pathway-level expression variation detects OY1 as a trans-eQTL hotspot affecting the expression of chlorophyll biosynthesis genes

To further explore the coordinated transcriptional regulation of the chlorophyll biosynthetic pathway by natural variation in maize, we identified 22 genes as chlorophyll pathway members (Figure 4, Supplemental Table S5). We calculated an index, as described above in our DEG analysis, using these 22 genes. This index is a summary statistic to estimate the aggregate effects of this pathway. The index values are the average of the z-score for the normalized counts from each transcript for every individual in the population sampled for eGWAS. We calculated the index using the chlorophyll biosynthesis genes for all three leaf tissues. Using the chlorophyll biosynthesis index as our trait, we performed GWA to identify SNPs associated with coordinated expression changes in the pathway (Supplemental Figure S8, Supplemental Table S13). We first explored the influence of the well-established polymorphism at *oy1* on the chlorophyll biosynthesis index. Significant SNP associations at the *oy1* locus were identified in L3Base and LMAD, indicating that the polymorphism at *oy1* influenced transcript accumulation in the chlorophyll biosynthesis pathway (Supplemental Figure S8 and Table 4). Three of five SNPs with effects on the chlorophyll biosynthetic index were also trans-eQTL, affecting the accumulation of transcripts encoded by *gun4*, *mpec1*, and *nyc1l1* (Table 2) and two (10-8968003 and 10-9004446) are displayed in Figure 6. Like the hot top SNPs and most hotspots, these SNPs did not have significant associations with mutant or wild type chlorophyll accumulation traits.

**Table 4.**
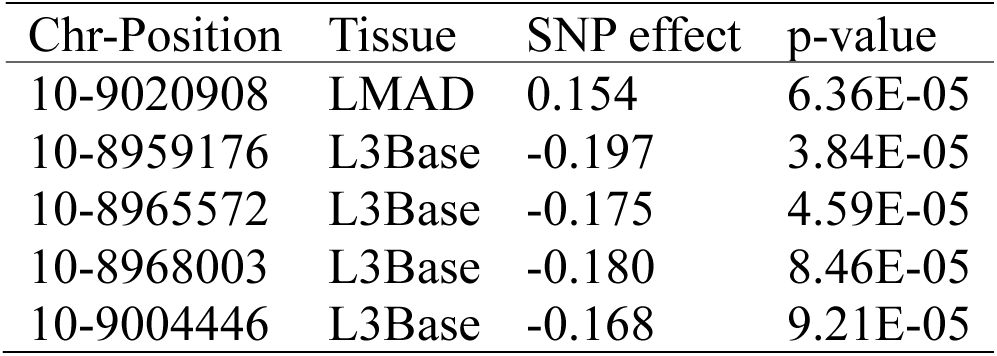
SNPs at the *oy1* locus associated with the chlorophyll biosynthesis index.

| Chr-Position | Tissue | SNP effect | p-value |
| --- | --- | --- | --- |
| 10-9020908 | LMAD | 0.154 | 6.36E-05 |
| 10-8959176 | L3Base | -0.197 | 3.84E-05 |
| 10-8965572 | L3Base | -0.175 | 4.59E-05 |
| 10-8968003 | L3Base | -0.180 | 8.46E-05 |
| 10-9004446 | L3Base | -0.168 | 9.21E-05 |

A genome-wide association using the chlorophyll biosynthesis index from each of the three tissues as traits identified eleven SNPs at p-value ≤ 10^-7^ (Table 5). The SNP 9-77570763 affecting the variation in the L3Base chlorophyll biosynthesis index was a GWAS hit and is within a Hotspot (Table 5). This SNP affects the expression of only four genes at the 10^-4^ threshold used for the hot SNP criteria, so does not appear as a hot SNP. Similarly, the SNPs detected in L3Base on chromosome 8 at 59Mb are within a hotspot but did not appear as hot SNPs (Table 5). All three index hits from LMAD also detected SNPs within hotspots (Table 5). Neither of the index hits on chromosome 2 identified hot SNPs, but the Index GWAS hit on chromosome 7 for the LMAD index was identified as a hot top SNP for gene expression in LMAD (Table 5). The index hit at 10-10925524 affecting the index in LMAN, was also within hotspots for LMAN trans-regulation of the pathway and was identified as a hot top SNP (Table 5). The Index hit on chromosome 2 at 29Mb was detected as a hot SNP but not as a hotspot (Table 5). Though not all loci identified this way overlapped with our hotspots and hot SNPs, this demonstrates that counting significant associations to identify hotspots/hot SNPs and the parametric sums of expression effects of the index calculation can arrive at the same loci.

**Table 5.** SNPs associated with the chlorophyll biosynthesis index at a p-value less than 10^-7^.

| Chr-position | SNP Effect | p-value | Trans regulatory hotspot class | Tissue |
| --- | --- | --- | --- | --- |
| 2-2949376 | -0.221 | 2.64E-08 | Hot SNP | LMAN |
| 2-149706803 | -0.220 | 7.43E-08 | Within Hotspot | LMAD |
| 2-183572149 | -0.242 | 1.19E-08 | Within Hotspot | LMAD |
| 7-13071884 | -0.257 | 9.72E-08 | Hot Top SNP | LMAD |
| 8-69497004 | 0.229 | 4.15E-08 | Within Hotspot | L3Base |
| 8-69506048 | 0.238 | 3.73E-08 | Within Hotspot | L3Base |
| 8-69509342 | 0.232 | 8.90E-08 | Within Hotspot | L3Base |
| 8-69547802 | 0.228 | 7.70E-08 | Within Hotspot | L3Base |
| 8-69571295 | 0.233 | 2.68E-08 | Within Hotspot | L3Base |
| 9-77570763 | -0.230 | 7.32E-08 | Within Hotspot | L3Base |
| 10-10925524 | -0.160 | 6.80E-08 | Hot Top SNP | LMAN |

### Does natural variation affecting expression level in tetrapyrrole pathway predict *Oy1-N1989* effects on plant growth and reproductivity maturity?

*Oy1-N1989/+* mutants show defects in growth and development, including delayed reproductive maturity, reduced stalk width, and altered plant height. These traits were strongly correlated with CCM values [26,27]. While *Oy1-N1989/oy1*^B73^:B73 mutants are taller than their wild-type siblings, *Oy1-N1989/oy1*^Mo17^:b094 mutants show a severe reduction in height [26,31]. To evaluate if morphological and growth defects in *Oy1-N1989/+* mutants were associated with changes in the expression of tetrapyrrole pathway genes, we intersected the SNPs encoding cis eQTLs (p-value <1e-4) with SNPs (p-value < 1e-4) linked to stalk width, plant height and flowering time in 343 F1 families [31]. We identified cis-acting SNPs linked to several tetrapyrrole pathway genes associated with time to reproductive maturity in *Oy1-N1989* mutants (Supplemental Figure S9, S10, S11). As expected from prior work [25–27,31], cis variation at the *oy1* locus in LMAN tissue was associated with mutant DTA and DTS, as well as difference DTA, difference DTS, and ratio DTS (Supplemental Figure S10, Supplemental Table S14). Reduced *oy1* expression delayed the days to reproductive maturity in the *Oy1-N1989/+* mutant and acted to enhance the mutant phenotype.

We also identified cis-acting alleles at *les22* that affected reproductive maturity. Increased accumulation of LES22 transcripts in LMAD and LMAN was associated with a delay in reproductive maturity in the mutants. This was affected by the same allele of *les22* that also influenced CCM values, indicating that the likely mechanism was a greater accumulation of chlorophyll. The allelic effect directions for DTS and DTA showed the same contrarian impact observed for CCM, where more transcripts from an upstream step resulted in more severe phenotypic effects in the *Oy1-N1989*/+ mutants. Cis variation affecting increased transcript accumulation from a second homolog of *les22, les22-like3* (*les22l3*; Zm00001d044186) also delayed mutant DTA and DTS (Supplemental Figure S9, S10, S11) recapitulating the contrarian effect direction at another of the paralogs encoding the conserved duplicates of this gene described above (Supplemental Figure S7). Cis-acting alleles associated with higher expression of a protoporphyrinogen oxidase, *prop3* (Zm00001d040539), in all leaf tissues were also significantly associated with a longer anthesis-silking interval in mutants. We also identified cis eQTLs associated with chlorophyll catabolism genes including *lethal leaf spot1* (*lls1*, Zm00001d027656)*, chlorophyllase2* (*chph2*, Zm00001d032926) and the chlorophyllide b reductase encoded by *non-yellow coloring1-like1 (nyc1l1*, Zm00001d013651) that had significant effect on reproductive maturity date in *Oy1-N1989/+* mutants (Supplemental Figure S9, S10, S11, Supplemental Table S14). This demonstrates that the impact of *Oy1-N1989* is sensitive to changes in the accumulation of transcripts encoding steps in chlorophyll biosynthesis and catabolism, as well as upstream steps in porphyrin biosynthesis.

Significant links were also detected between gene expression and variation in stalk width. As demonstrated previously [31], cis variants linked to higher expression of *oy1* increased the ratio of mutant to wild type stalk width and decreased the difference between wild type and mutant stalk width. In the eGWAS analysis, SNPs affecting *oy1* expression in LMAN were significantly associated with stalk width (Supplemental Figure S10, Supplemental Table S14). Cis SNPs associated with the expression of *pcr1* also affected the ratio and difference of stalk widths between *Oy1-N1989*/+ and wild-type F1 siblings (Supplemental Figure S9, S10, S11). The SNP effect direction on gene expression in L3Base, LMAN, and LMAN was consistent with reduced gene expression of *pcr1,* leading to a reduction in mutant stalk width. Once again, the effect of variation in *les22* on phenotypic expression was opposite to that of the other porphyrin pathway genes. SNPs affecting a decrease in expression of *les22* in mature leaf tissues were associated with increased stalk width in the mutant. Of all the cis SNPs linked expression, only those at *elm2* in LMAN were significantly associated with wild type stalk width (Supplemental Table S14), consistent with repeated observations of the effects at this locus and trans regulation of this heme oxygenase affecting phenotypic traits and regulation (Supplemental Tables S10, S11, and S12).

We also identified significant associations of cis-regulatory variation at tetrapyrrole pathway genes with mutant height. We did not observe any association between the accumulation of OY1 transcripts and either mutant or wild type height traits, as expected for the complex QTL effects at this locus [31]. Among the other genes in the chlorophyll biosynthesis pathway, only the SNPs associated with higher expression of Mg-chelatase subunit H (*chlh1*) in L3Base were linked to a reduced ratio of mutant to wild type flag leaf height (Supplemental Table S14, Supplemental Figure S11). No cis eQTLs at other core chlorophyll biosynthetic genes had significant associations with height traits. However, cis variants for genes in the porphyrin biosynthesis pathway were found to be significantly associated with height traits in *Oy1-N9189/+* mutants. The cis SNPs associated with *glutamyl tRNA synthetase2* (*gts2*, Zm00001d015037) in L3Base and LMAN had significant effects on the ratio of wild type to mutant and the difference between wild type and mutant height traits (Supplemental Table S14, Supplemental Figure S10, Supplemental Figure S11). Like the observation for reproductive maturity, reduced expression of *prpo3* in all leaf tissues was significantly associated with an increased mutant ear-to-flag leaf height. Reduced expression of *prpo1* in the three leaf tissues was linked to decreased flag leaf height in both wild-type plants and mutants, as well as mutant ear height. The inconsistent effect directions for SNPs linked to *prpo1* and *prpo3* and the effect on both mutant and wildtype traits at *prpo1* suggest these are not changes affected by chlorophyll levels. Several other cis variants were associated with differences in wild type height traits but not mutant traits, indicating they were not rate-limiting for *Oy1-N1989/+* phenotype expression and may not be due to changes in chlorophyll accumulation (Supplemental Table S14).

Among the chlorophyll-degrading genes, we identified multiple associations that affected mutant heights. These included 24 cis SNPs associated with *lls1* expression in L3Base with a significant effect on mutant flag leaf and ear height. Of these, three were significantly associated with increased wild type plant height. A cis polymorphism linked to reduced expression of *lethal leaf spot1 like5* (*lls1l5*, Zm00001d011532), a homolog of *lls1,* in LMAN and LMAD also reduced the difference between wildtype and mutant ear height. Cis SNPs linked to the expression of *nycl1* had similar effects on both mutant and wild type flag leaf height (Supplemental Figure S9, S10, S11, Supplemental Table S14). As these did not also affect changes in CCM values, it is unclear if this results from a failure to detect CCM GWAS associations or some other cause.

We also tested the association of morphological traits with SNPs associated with other branches of the porphyrin pathway, including heme, siroheme, and bilin pathways. Increased expression of *fc2* (Zm00001d016854) in LMAD and LMAN was associated with higher flag leaf height and ear height ratios. However, the relationship between SNP effects on gene expression and height was opposite in L3Base. The lower expression of *elm2* was associated with increased flag leaf height and ear height in both mutant and wild-type plants and increased ear-to-flag leaf height in wild-type plants. These effects in the younger leaves were consistent with the expectation for reduced phytochrome signaling.

### Expression data illuminate possible causes of the morphological and growth effects of *Oy1-N1989*/+ mutants

The defects in plant growth and development in *Oy1-N1989/+* mutants respond to the natural allelic variation in *oy1* in a complex manner [25–27,31]. We performed a careful pathway-level and candidate gene analysis of the RNA-seq data to search for the mechanisms underlying the morphological variation in *Oy1-N1989*/+ mutants. *Oy1-N1989* suppresses tillering in maize mutants, including exhibiting an epistatic interaction with mutants in *teosinte branched1 (tb1,* Zm00001d033673*), grassy tillers1 (gt1*, Zm00001d028129), and sweet corn tillering that includes effects of *tiller number1* (*tin1*, Zm00001d018816) [26]. We examined the expression of known tillering regulators in our RNA-seq of *Oy1-N1989/+* mutants. Transcripts from *tin1*, known to increase tillering when overexpressed, had significantly lower expression in the enhanced *Oy1-N1989/oy1*^Mo17^:b094 mutant (Supplemental Table S1). Previous work demonstrated that *oy1* was epistatic to the cis-regulatory eQTL resulting from the intron retention allele of *tin1* in the Purdue sweetcorn line P51 [23], suggesting that *oy1* mediates decreased tillering via this gene [26]. Neither *gt1* nor *tb1* transcripts accumulated in our RNA-seq data from leaf tissue. We also analyzed the effects of *Oy1-N1989/+* on transcripts encoding strigolactone biosynthesis, a hormone that suppresses tiller bud outgrowth [38–40]. Transcripts of two maize homologs of *dwarf27* (*d27*) encoding a beta carotene isomerase, *dwarf27-like1* (*d27l1*, Zm00001d007560) and *dwarf27-like2* (*d27l2,* Zm00001d007549), were reduced in *Oy1-N1989/+* mutants. Transcripts of gene encoding the next step in strigolactone biosynthesis, *carotenoid cleavage dioxygenase7* (*ccd7*, Zm00001d002736), were significantly reduced in both *Oy1-N1989/oy1*^B73^ and *Oy1-N1989/oy1*^Mo17^:b094 mutants (Supplemental Table S15). Further studies are required to identify if there is reduced strigolactone biosynthesis or if the reduced expression of strigolactone biosynthetic genes results from feedback regulation.

The effects of *Oy1-N1989/+* on plant height are complex and incompatible with simple physiological explanations, such as a reduction in photosynthate simply reducing growth. In our previous study, modifying mutant severity by genetic background reverses the effect of *Oy1-N1989/+* on plant height [26]. When the suppressing *oy1*^B73^ is present, *Oy1-N1989/oy1*^B73^:B73 is taller than its wild-type siblings. However, when the enhancing *oy1*^Mo17^ allele is present, *Oy1-N1989/oy1*^Mo17^:b094 is shorter than the wild type siblings [26]. Therefore, the fold reductions in chlorophyll accumulation in *Oy1-N1989*/*oy1*^B73^ increased plant height, but a further drop in chlorophyll in *Oy1-N1989/ oy1*^Mo17^ resulted in short-statured plants relative to the wildtype siblings. This phenotype was observed in every population that segregated for the modifier alleles at *oy1* evaluated to date (tens of thousands of plants; [25,26,31]). To seek molecular insight into this phenotype, we analyzed our RNA-seq experiments for the gene expression effects of *Oy1-N1989/+* mutants on pathways that affect plant height, including gibberellin (GA) biosynthesis, GA signaling, sterol and brassinosteroid (BR) biosynthesis, and BR signaling (Figure 9, Supplemental Table S15). Among the GA biosynthesis and signaling pathway genes, only transcripts of *gibberellin 20-oxidase3* (*ga20ox3,* Zm00001d042611) were significantly reduced at the experiment-wide false positive corrected threshold in both the enhanced and suppressed mutants. In the enhanced *Oy1-N1989/oy1*^Mo17^:b094 background, transcripts of *gibberellin 2-oxidase13* (*ga2ox13*, Zm00001d039394), *dwarf8 (d8,* Zm00001d033680), and *dwarf9 (d9,* Zm00001d013465) were also reduced significantly. The transcripts of *MYB-transcription factor 74* (*myb74*, Zm00001d012544) were significantly increased at the experiment-wide false positive corrected threshold in *Oy1-N1989* compared to wild-type siblings. No BR or sterol biosynthetic genes or BR signaling genes were differentially expressed at the experiment wide-false positive corrected threshold in the suppressed *Oy1-N1989/oy1*^B73^:B73 mutants. However, eleven genes were DEG in the enhanced and shorter *Oy1-N1989/oy1*^Mo17^:b094 mutants. These included steps in sterol biosynthesis that may respond to lower growth and three of the P450s encoding steps in BR biosynthesis. However, the expression directions were mixed with some transcripts increasing and others decreasing in accumulation (Figure 9; Supplemental Table S15). During the preparation of this manuscript, a new report demonstrating that ABCB19 is a BR transporter was published [41]. The transcript encoded by Zm00001d025590, one of three ABCB19 paralogs in maize, increased accumulation in the shorter *Oy1-N1989/oy1^Mo17^*:b094 mutants. None of these results provides a clear mechanism for the increased plant height in *Oy1-N1989/oy1*^B73^:B73 mutants but were consistent with loss of BR and sterols in shorter enhanced *Oy1-N1989/oy1^Mo17^*:b094 mutants.

**Figure 9.**
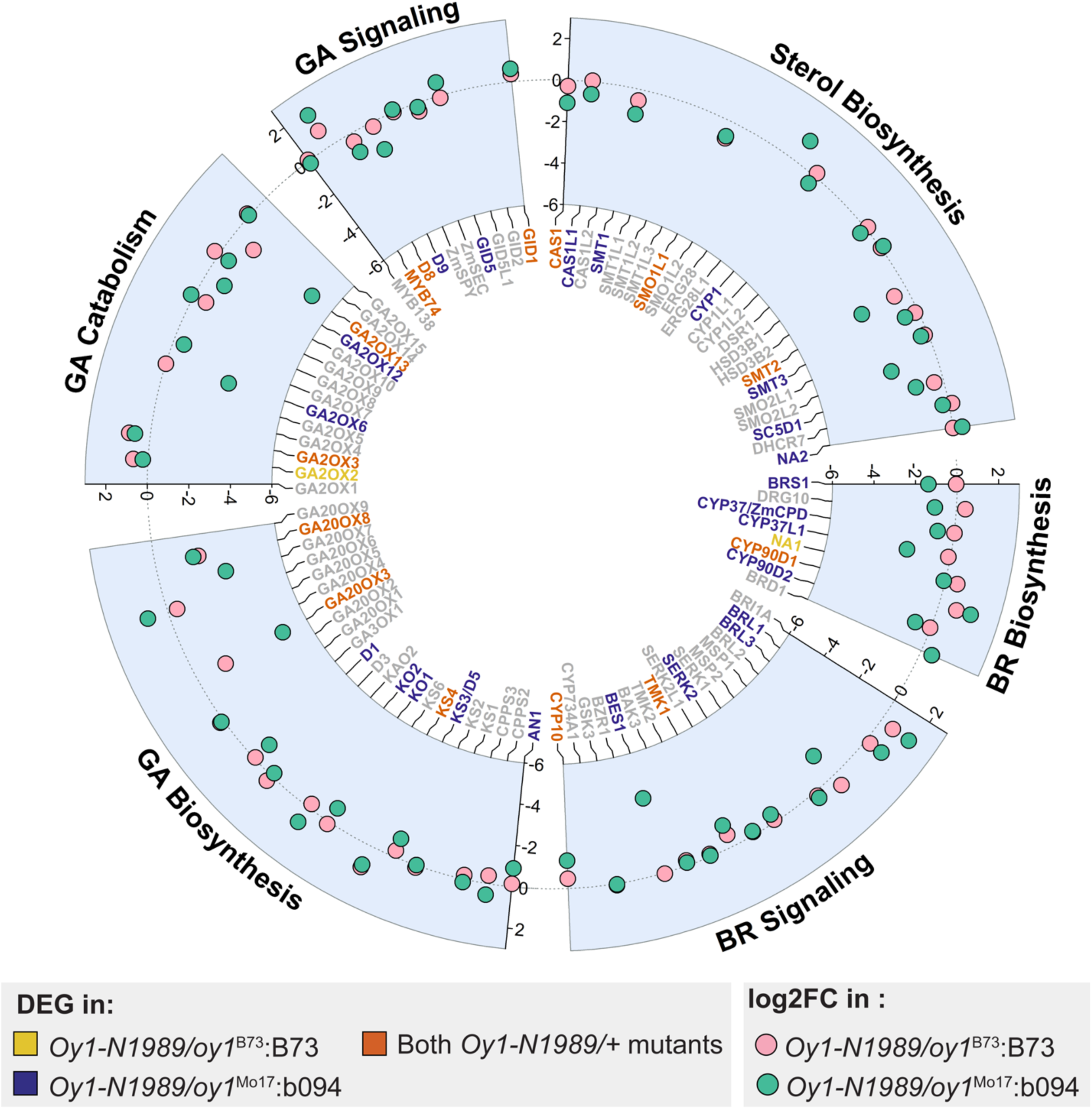
Circle plot showing log2 fold change of transcripts of genes encoding steps in gibberellin (GA) and brassinosteroid (BR) biosynthetic and signaling pathways in *Oy1-N1989/oy1*^B73^:B73 (pink circles) and *Oy1-N1989/oy1*^Mo17^:b094 (green circles) mutants as compared to their respective wild-type controls. The transcripts highlighted in red were differentially expressed in both *Oy1-N1989/+* mutants at unadjusted p-value (p ≤ 0.05). Transcripts highlighted in yellow and dark blue were only differentially expressed in Oy1-N1989/oy1^B73^:B73 and *Oy1-N1989/oy1*^Mo17^:b094 respectively. Transcripts in grey were either not differentially expressed at unadjusted p-value cutoff of 0.05 or were not detected in our RNA-seq data.

We evaluated the impact of biosynthesis and signaling of gibberellic acid, sterol, brassinosteroids, and auxin pathways more holistically by using the less stringent criteria of p-value ≤ 0.05 to identify gene expression patterns at the pathway level. Transcripts from five of the eight BR biosynthetic steps were decreased at this threshold in the enhanced *Oy1-N1989/oy1^Mo17^*:b094 mutants (Figure 9). The possibility of BR involvement in the height differences between enhanced and suppressed *Oy1-N9189/+* mutants was highlighted by the direction of gene expression affected in the background-enhanced mutants. For example, the rate-limiting BR biosynthetic genes encoding the maize homologs of Arabidopsis *constitutive photomorphogenic dwarf/dwarf3/cabbage* (*cyp37*), *cyp90D* (*cyp90d2*), and *dwf4* (*brs1*) were significantly decreased in accumulation in the enhanced *Oy1-N9189/+* mutants but not affected in the suppressed *Oy1-N1989/oy1*^B73^:B73 mutants (Figure 9, Supplemental Table S15). Among the 15 sterol biosynthetic genes expressed in the two mutants, eight showed reduced transcript accumulation in *Oy1-N1989/oy1*^Mo17^:b094 (Figure 9). In addition to effects at *abcb19*, transcripts encoded by *brachytic2* were unaffected in the suppressing background mutants and decreased in accumulation in the shorter enhanced mutants. Transcripts of genes encoding steps in GA biosynthesis in maize including *anther ear 1* (*an1,* Zm00001d032961), *dwarf5* (*d5,* Zm00001d002349), and *dwarf1* (*d1,* Zm00001d039634), were also reduced in the enhanced *Oy1-N1989/oy1^Mo17^*:b094 mutants but not altered in *Oy1-N1989/oy1*^B73^:B73. In addition to *ga20ox3* (Zm00001d042611), transcripts of *ga20ox8* (Zm00001d049926) were decreased in both mutants*. Dwarf8* (*d8,* Zm00001d033680), a key repressor of GA signaling in maize, was repressed in both *Oy1-N1989/oy1*^B73^:B73 and *Oy1-N1989/oy1*^Mo17^:b094 mutants with higher reduction in *Oy1-N1989/oy1*^Mo17^:b094. Transcripts of GA receptor *gid1* were increased in *Oy1-N1989/+* mutants in both genetic backgrounds. Enhanced mutant *Oy1-N1989/oy1*^Mo17^:b094 also accumulated transcripts of a GA signaling gene, *gid5* (Zm00001d016973). The reduction in the accumulation of transcripts encoding a GA signal repressor and the increased accumulation of the GA receptor (Figure 9) may explain why *Oy1-N1989/oy1*^B73^:B73 is taller than wild-type siblings. The more dramatic reduction of GA biosynthesis in *Oy1-N1989/oy1*^Mo17^:b094 (Figure 9) may explain the reversal of this effect in *Oy1-N1989/oy1*^Mo17^:b094.

Auxin biosynthesis and signal transduction pathway also displayed altered expression in *Oy1-N19*89/+ mutants (p-value ≤ 0.05). The *Oy1-N1989/+* mutants showed reduced accumulation of transcripts encoding the auxin biosynthetic gene, *vanishing tassel2* (*vt2*, Zm00001d008700) (Supplemental Table S15). Transcripts of the gene encoding the auxin transporter *pin-formed protein2* (*pin2*, Zm00001d018024) were decreased in both *Oy1-N1989/+* mutants. The genetic backgrounds had opposing effects on the expression of *pin-formed protein10* (*pin10*, Zm00001d044083), where the tall *Oy1-N1989/oy1*^B73^:B73 plants had reduced accumulation of PIN10 transcripts and the shorter *Oy1-N1989/oy1*^Mo17^:b094 plants showed higher PIN10 transcripts as compared to their respective wild-type siblings. *Oy1-N1989/oy1*^Mo17^:b094 also showed increased accumulation of transcripts of *pin-formed protein3* (*pin3*, Zm00001d052269), which was not significantly affected in *Oy1-N1989/oy1*^B73^:B73. Transcripts of *transport inhibitor response1* (*tir1*, Zm00001d010863) encoding an auxin receptor were significantly decreased in *Oy1-N1989/+* mutants in both genetic backgrounds. Of 33 auxin response factor (ARF) family genes, transcripts of 14 genes were significantly affected in *Oy1-N1989/oy1*^Mo17^:b094 (Supplemental Table S15). Transcripts of 12 of these 14 genes were reduced in accumulation in *Oy1-N1989/oy1*^Mo17^:b094 mutant. Of the 33 *arf* family genes, only four were significantly affected in *Oy1-N1989/oy1*^B73^:B73 with no clear direction of effect on gene expression in the mutant compared to its congenic wild type, and no clear indication from the gene expression patterns what if any effect auxin has on this unusual relationship between chlorophyll biogenesis and plant height. This remains a mystery deserving an alternative experimental approach.

## Discussion

### The wild-type allele at *oy1* determines the severity of gene expression consequences in *Oy1-N1989/+* mutant

In this study, we examined the transcriptional consequences of variation in the *oy1* gene encoding subunit I of the Mg-chelatase enzyme in maize. We used the semi-dominant *Oy1-N1989* dominant-negative mutant allele in heterozygous plants with differing phenotypic severity conditioned by the wild-type allele at the *oy1* locus. Consistent with the impact on morphology, *Oy1-N1989/+* mutants carrying the enhancing *oy1*^Mo17^ allele showed a more severe effect on gene expression than mutants carrying the suppressing *oy1*^B73^ allele. This was evident from the number of DEGs in RNA-seq analysis of transcript abundances in the two individuals as compared to their congenic wild types, where the enhanced *Oy1-N1989/oy1*^Mo17^:b094 affected many more DEGs than *Oy1-N1989/oy1*^B73^:B73 (Figure 2). Among the DEGs affected in both genetic backgrounds, the transcript levels of the shared genes were impacted in the same direction by *Oy1-N1989* irrespective of the wild-type allele at *oy1*. The clearest demonstration that *oy1*^Mo17^ enhances the severity of the mutant phenotype and does not have a separate effect came from the analysis of the magnitude of effects on gene expression. The enhanced phenotype of *Oy1-N1989/oy1*^Mo17^:b094 resulted in a greater fold-change in the expression of genes that were DEGs due to *Oy1-N1989/+* in both backgrounds (Figure 2). This was further demonstrated using expression indices that measure the aggregate effect on gene expression in the two mutants and wild-type samples. Transcripts induced by *Oy1-N1989/oy1*^B73^:B73 were accumulated to a greater extent in *Oy1-N1989/oy1*^Mo17^:b094 (Figure 3). Greater changes in transcript accumulation were observed in the enhanced mutants regardless of the gene sets tested. These results indicate that genetic background affected by the *oy1* locus determined the severity of gene expression effects due to *Oy1-N1989*. The use of an expression index to look at the coordinate regulation of gene expression was validated by the fact that transcriptional hotspots colocalized with GWAS peaks obtained using index values calculated from expression data from an association mapping panel (Table 5).

### Variation in gene expression of *oy1* wild-type alleles affects the severity of the *Oy1-N1989*/+ mutant phenotype

We previously demonstrated that cis-regulatory polymorphism at *oy1* is associated with a greater accumulation of OY1 transcripts from the B73 allele using NILs, RILs, and F1 hybrids [25]. Lower expression from wild type *oy1* increased the phenotypic severity of *Oy1-N1989/+* on chlorophyll accumulation [25]. We compared the cis variation at *oy1* in diverse maize lines with the SNPs affecting chlorophyll accumulation in mutant F1 hybrids. All the cis-acting SNPs associated with expression variation in *oy1* in our two mature leaf tissues (LMAD and LMAN) had significant associations with mutant chlorophyll traits (Figure 5). The cis-regulatory alleles that increased the expression of *oy1* were consistently associated with higher chlorophyll contents in the mutants. Thus, lines with higher expression of wild-type allele might produce higher amounts of wild-type gene product that competes with the mutant gene product, thereby suppressing the phenotype in the mutants.

### Loss of Mg-chelatase activity affects the expression of tetrapyrrole pathway genes

In plants, all tetrapyrroles are synthesized from a glutamyl-tRNA precursor in a multi-branched pathway [2,4,42]. In this study, we identified 70 maize genes encoding steps in all the branches of the tetrapyrrole pathway. (Figure 1, Supplemental Table S5). We detected pathway-level effects by bringing these gene annotations into our RNA-seq analysis and observed coordinated regulation of successive steps and branches (Figure 4). *Oy1-N1989*/+ plants displayed increased transcript levels for genes encoding the steps of the pathway up to protoporphyrin IX biosynthesis (Figure 1 and Figure 4). This demonstrates the existence of a transcriptional regulation of the pathway that affects more than just glutamyl tRNA reductase in response to a mutation of Mg-chelatase. The altered transcript levels of upstream ALA and porphyrin biosynthetic genes suggest that a transcriptional mechanism either senses Mg-chelatase activity or responds to the accumulation of biosynthetic intermediates. The accumulation of some intermediates between ALA and protoporphyrin IX in the presence of light and oxygen damages the cell [36]. If these intermediates had accumulated in the mutant, we would have seen phototoxic damage phenotypes in *Oy1-N1989/+* mutants, but we did not. The lack of such phenotypes suggests that the increased transcription from these genes did not cause the accumulation of toxic intermediates in the mutant. The reduced transcript abundance of *sum1* (Figure 4, Supplemental Table S5), which encodes the first committed step in the siroheme branch, suggests a compensatory regulatory mechanism may exist to reduce flux in the siroheme pathway during a loss of chlorophyll. Other genes in this branch did not show a similar decrease in expression. We did not observe transcriptional regulation of the heme branch as the transcript levels of genes encoding steps in the heme and phytochromobilin branch were not affected in *Oy1-N1989*/+ mutants.

*Oy1-N1989/+* altered the expression of genes encoding steps in the chlorophyll biosynthesis branch. In both genetic backgrounds, *Oy1-N1989/+* increased CHLD transcripts, demonstrating a compensatory feedback regulation at the transcription of Mg-chelatase. Prior work with a rice loss-of-function allele in the *OsCHLI* gene resulted in similar gene expression patterns with increased accumulation of upstream porphyrin biosynthesis steps, increased CHLD, and decreased GUN4 transcripts [43]. This similarity provides further confirmation of the dominant-negative nature of the *Oy1-N1989* allele. The consequences of *Oy1-N1989* and its modifiers on the transcript levels of genes encoding other enzymes were more complicated. When the *oy1^B73^* allele was present, transcript levels of CHLH, a subunit of Mg-chelatase, showed a compensatory increase. But when the *oy1^Mo17^* allele was present, the enhanced mutant phenotype included decreased transcript levels of CHLH, raising the possibility that one contribution to the severe phenotype of *oy1^Mo17^* enhanced *Oy1-N1989/+* mutants was reduced transcription from genes encoding steps in the chlorophyll biosynthetic pathway rather than a compensatory upregulation. The *oy1^Mo17^* enhanced *Oy1-N1989/+* mutants also had significantly fewer transcripts from genes in the chlorophyll cycle (Figures 1 and 2; Supplemental Table S1). This suggests a transcriptional response to severe reduction in chlorophyll accumulation as these steps would require less flux when less metabolite was present. This demonstrates yet another detection of a set of co-regulated genes affected by an understudied transcriptional regulation in this pathway. Similarly, transcripts of genes encoding steps in chlorophyll catabolism, including *nye1*, *nye2,* and *rccr1,* were all reduced in accumulation in *Oy1-N1989/+* mutants, suggesting a decrease in chlorophyll catabolism when chlorophyll was limited and demonstrating a transcriptional regulatory mechanism acting at the end of the pathway as well. Once again, exploring the gene expression consequences of metabolic mutants highlights regulatory pathways not previously detected.

### Natural variation affected transcriptional co-regulation of genes encoding steps in tetrapyrrole biosynthesis

One disadvantage to using mutants is the outsized impact they often have on metabolism and the potential for complex secondary effects. Death, for example, is the ultimate pleiotropic phenotype. By contrast, metabolic feedback in biochemical pathways is constructed during evolution in response to a narrower physiological range of gene products (transcripts and protein) and metabolite levels. One way to avoid the pitfalls of overinterpreting mutant phenotypes and to seek physiologically relevant regulation is to explore natural variation. In this study, eGWAS of 70 genes in the tetrapyrrole pathway in wild-type maize lines identified abundant cis and trans regulation affecting the of these genes in three leaf tissues. Remarkably, SNPs linked to the *oy1* locus were identified as trans-regulators of the pathway. This included SNPs affecting the expression of 25 genes in LMAD, 17 in LMAN, and 28 in L3Base at the permissive 10^-4^ p-value cutoff (Supplemental Table S8). This suggests that the feedback affected by the *Oy1-N1989* allele is not an artifact of an unusual mutant allele and can also be affected by natural variation at this locus. However, it remains formally possible that another linked gene affects transcriptional regulation of the tetrapyrrole pathway. It demonstrates that these genes are co-regulated and that the natural variants are of sufficient consequence to cause compensatory transcriptional regulation of genes encoding steps in the tetrapyrrole pathway. This was further demonstrated through GWAS using gene expression index value derived from 22 genes in the chlorophyll biosynthesis and chlorophyll cycle. This aggregate trait, called the chlorophyll index, was also affected by SNPs linked to *oy1* (Table 4). Together with the RNA-seq data from *Oy1-N1989/+* mutants in enhancing and suppressing genetic backgrounds, these data indicate that variation in *oy1* triggers a transcriptional checkpoint regulating the tetrapyrrole biosynthetic pathway.

To further explore the transcriptional regulation of the tetrapyrrole biosynthetic pathway, we identified top SNPs that affected the expression of eight or more genes in trans. These hot top SNPs permit testing of the direction of expression effects for each allele. The analysis of the genes affected by 55 SNPs that were the top SNP association for eight or more genes showed that transcript levels of nearly all genes affected by these trans-regulatory hotspot alleles were co-regulated (Figure 7). These genes encode the initial common steps of the tetrapyrrole pathway and steps in the chlorophyll branch. Like the coordinate direction of expression effects (Figure 7), a subset of steps in the porphyrin and chlorophyll pathway were more commonly affected by transcriptional hotspots than any other steps of the pathway when assessed either as windowed hotspot loci or as single hot top SNPs (Figures 7 and 8). The two mature leaf tissues tested here identified an order of magnitude fewer hot top SNPs with only two hot top SNPs in LMAD and three in LMAN.

Only one gene displayed divergent trans-regulatory consequences in the hot top SNPs: *cf1*. This gene is only divergently affected in expression level in mature leaves during the day. It may be that the different regulation in the leaf base and the mature leaf underlies the unusual phenotype expression. The SNPs associated with trans hotspots in LMAD had the opposite effect on the expression of *cf1* compared to other co-regulated porphyrin and chlorophyll biosynthetic genes in the pathway. In contrast, transcript levels of *cf1* were coordinately regulated with transcripts of porphyrin and chlorophyll biosynthetic genes in LMAN and L3Base. The gene *cf1* encodes a porphobilinogen deaminase that catalyzes the conversion of porphobilinogen to hydroxymethylbilane, the precursor of uroporphyrinogen III. Hydroxymethylbilane is unstable and can spontaneously cyclize to form photoreactive uroporphyrinogen I. The *cf1* gene might be differently regulated in mature leaf tissues to prevent the accumulation of phototoxic intermediates and thus prevent photodamage. One could argue that if there is a negative regulation of *cf1* during the day to reduce the accumulation of phototoxic intermediates, we should see similar effects in the base of third leaf, which was also collected during the day. However, we do not see opposing regulation of *cf1* in this tissue. This suggests that regulation of the pathway is dependent on the developmental stage. Perhaps consonant with this, both the weak alleles found in *cf1* mutants and the strong loss of function allele encoded by *necrotic3* result in diurnal bands of brown tissue in seedlings [19,44]. The upregulation of enzymes upstream in porphyrin metabolism by *Oy1-N1989* (Figure 4, Supplemental Table S5) and feedback regulation of *cf1* by alleles of *oy1* (Supplemental Table S8) might explain the suppression of *cf1-m2* by *Oy1-N700*. The *cf1-m2* allele is encoded by a mu insertion in the 5’UTR limiting transcription [19] as are all four camouflage alleles. In this light, the fact that *oy1* in wild-type maize acted as a trans regulator of *cf1* and that *cf1* showed changes in expression in *Oy1-N1989/+* mutants may explain the paradoxical suppression of the mutant phenotype in *cf1* by the mutation of the downstream *Oy1-N700/+*.

### Natural variation in the expression of tetrapyrrole pathway genes impacts the severity of *Oy1-N1989* phenotype

Investigation of natural variation affecting the expression of tetrapyrrole metabolism genes identified cis-regulatory variation linked to these genes (Supplemental Figure S3). As described above, SNPs affecting cis-acting regulation of the *oy1* locus in mature leaf tissues also influenced the chlorophyll contents of the mutants in F1 association mapping GWAS experiments (Figure 5). All cis variants associated with *oy1* expression, identified using RNA extracted from wild-type inbreds in a prior study [30], were also associated with variation in mutant chlorophyll contents, but not wild type chlorophyll contents from our previous F1 Association mapping experiments (Figure 5; [25]). The joint performance of these SNPs across two different experiments represents true biological replicates and even utilizes inbreds and F1 hybrids as the basis of the material. We explored if SNPs affecting cis-regulatory expression variation at other tetrapyrrole pathway genes also affected the mutant chlorophyll content in the F1-association mapping experiment. By cross-tabulating the impact of the cis variants at all genes on chlorophyll content in both the WT and *Oy1-N1989* mutant F1 hybrids, we identified additional natural variants impacting both the gene expression and chlorophyll content (Supplemental Figure S3, Supplemental Table S7). However, unlike the results obtained for *oy1*, not all cis variants associated with the expression of other tetrapyrrole pathway genes showed an impact on chlorophyll content even at a suggestive p-value cut-off of 1e-4.

### Transcriptional hotspot analyses rely on a false and dangerous assumption

All assessments of hotspots, here and elsewhere, use a null hypothesis that relies on an assumption that gene expression is independent in each sample. While convenient and necessary for these calculations, this is naïve and false and will be trivially rejected in any experiment where the sample size is large enough. The tens of thousands of gene expression estimates from each sample share an abiotic environment, sampling time, developmental stage, wounding history, and biotic influence and so covariance in gene expression will be greater within a sample than between neighbors. An assumption of independence short-hands all causes of expression correspondence, and any fortuitous linkage with SNP variation, as genetic causation. As a result, we expect all hotspot calculations to overestimate the number of hotspots by whatever degree of inflation is caused by experimental design flaws and stochastic alignment of SNP variation and environmental features.

### Pathway-level analysis of RNA-seq data reveals possible causes of phenotypic effects of *Oy1-N1989*

The *Oy1-N1989* mutants show several defects in growth and development. The impact of *Oy1-N1989* on the height of heterozygous plants is variable depending on the genotype of the modifier encoded by the wild-type copy of *oy1*. *Oy1-N1989/+* mutants with the *oy1*^Mo17^ allele are reduced in height, whereas mutants in the B73 background are taller than isogenic wild-type siblings. This variation in height in *Oy1-N1989/+* mutants cannot be explained by a linear effect of the loss of chlorophyll biogenesis as reduced chlorophyll in *Oy1-N1989/oy1^B73^* as compared to wild-type controls results in taller plants while reduced chlorophyll in *Oy1-N1989/oy1^Mo17^* results in shorter plants than its wild-type controls. As expected for a non-linear relationship, the cis variation at *oy1* did not show an association with plant height traits, indicating that the effects on height are not direct consequences of defects in chlorophyll biogenesis but indirect effects of changes in expression of other genes or pathways. We carefully analyzed the genes encoding the biosynthesis and signaling components of two phytohormones in the RNA-seq data from *Oy1-N1989/+*. This pathway-targeted analysis of RNA-seq expression data revealed that our suppressed and enhanced *Oy1-N1989* mutants showed variable effects on genes in both GA and BR biosynthesis. Transcripts of biosynthetic genes in both GA and BR pathways were reduced in accumulation in enhanced *Oy1-N1989/+* mutants, suggesting that the reduced height of these mutants co-occurred with reduced biosynthesis of these phytohormones. However, the increased height of suppressed *Oy1-N1989/oy1^B73^* mutants was not accompanied by increased transcripts of genes encoding steps in GA or genes encoding rate-limiting steps in BR biosynthesis. Repressors of GA signaling, *d8,* and *d9* DELLA domain GRAS transcription factors, showed reduced accumulation, while the transcripts encoding GA receptors increased in their accumulation. We speculate that GA sensitivity could contribute to the height increases in these mutants. We propose that future experiments seek to untangle the surprising effect of the *Oy1-N1989/oy1^B73^* test for an effect of GA signaling.

## Material and methods

### Plant material

The mutant allele *Oy1-N1989* was obtained from maize genetics COOP and backcrossed for eight generations with B73, as described previously [25]. Pollen from the *Oy1-N1989/oy1* mutant introgressed into the B73 inbred background (*Oy1-N1989/oy1*^B73^:B73) was crossed to B73 to generate F_1_ hybrids segregating 1:1 for mutant and wild-type siblings. The mutant *Oy1-N1989/oy1*^B73^:B73 was also crossed to a near-isogenic line, b094, with introgression of the Mo17 allele at *oy1* (*oy1*^Mo17^) in a B73 background [25,27]. The progeny of this cross resulted in ∼1:1 segregation of mutant (*Oy1-N1989/oy1*^Mo17^:b094) and wild-type plants (*oy1*^B73^*/oy1*^Mo17^:b094) in a B73/b094 hybrid background [27]. Plants were grown at the Purdue Agronomy Center for Research and Education (ACRE) in West Lafayette, Indiana, during the summer of 2017. Standard fertilization, weed suppression, and pest control practices for maize cultivation were followed.

### RNA-sequencing and transcriptomic analysis

We collected the third leaf from field-grown plants at the V3 developmental stage. Six individuals of each genotype were pooled to make one biological replicate. Three biological replicates were collected from *Oy1-N1989/oy1*^B73^:B73 mutants, *Oy1-N1989/oy1*^Mo17^:b094 mutants, and their congenic wild-type siblings. Total RNA was extracted using TRIzol reagent (Invitrogen, CA, USA). The concentration and integrity of RNA were assessed using a Nanodrop 2000c spectrophotometer (Thermo Scientific, Waltham, MA). Novogene (Sacramento, CA) performed library construction and sequencing using the NovaSeq 6000 sequencing platform (Illumina, San Diego, CA).

Sequenced reads were aligned to the maize reference genome (B73 RefGen_v4, [45]) downloaded from MaizeGDB (https://download.maizegdb.org/Genomes/B73/Zm-B73-REFERENCE-GRAMENE-4.0/). Alignment was done using *bowtie2 v2.2.8* [46] and the gene models from the maize reference genome version 4 annotation were used to identify splice junctions. The number of reads aligned to each gene was used to derive counts using *htseq-count* [47]. Differential expression analysis was performed using DEseq2 v1.26.0 [48]. Transcripts with |log_2_ fold change| ≥ 1 and FDR adjusted p-value ≤ 0.05 were considered as differentially expressed (Supplemental Table S1). Unfiltered DESeq2 outputs are in Supplemental Table S2. Heatmaps were generated using the *pheatmap* package in R [49].

To calculate the index value for *Oy1-N1989/oy1* mutants and their congenic wild types, we derived Z-scores for the accumulation of each gene in the differentially expressed gene sets. The Z-score of each gene per sample was calculated from the normalized transcript counts as follows:

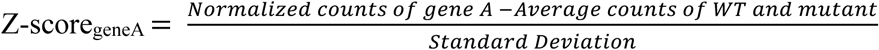

The index value for each sample was obtained by averaging the Z-scores for a set of genes identified as differentially expressed.

Gene ontology enrichment analysis for DEGs was performed using a web-based server *AGRIGO v2.0* ([50] using the following parameters: Fisher’s test, with Yekutieli (FDR under dependence) multi-test adjustment [51]. Plant GO Slim was selected as the gene ontology type. GO terms with FDR< 0.05 were selected as significant. Data were visualized using the *GOplot* package in R [52]. Z-scores for each GO term in the GOplot were calculated as follows:

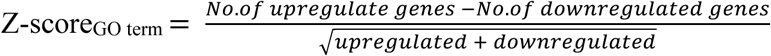

### Identification of genes involved in tetrapyrrole biosynthesis

We retrieved the genes involved in the tetrapyrrole biosynthesis pathway in Arabidopsis and maize from the KEGG pathway database [53,54] We obtained the orthologs of Arabidopsis genes in maize from PLAZA Monocots 5.0 [55]. We retrieved additional genes involved in the tetrapyrrole pathway in maize from the CornCyc database in the Plant Metabolic Network (PMN) data portal (https://pmn.plantcyc.org). The complete list of genes and their correct annotations and nomenclature in maize is provided in supplemental Table S5.

### Natural variation in transcript abundances

The Box-Cox transformed expression counts were obtained from a previously published study of RNA-sequencing data from 296 diverse maize lines [30]. We examined the seventy tetrapyrrole biosynthesis genes expressed in the following three leaf tissues: the base of the third leaf (L3Base), mature leaves collected during the day (LMAD), and mature leaves collected during the night (LMAN).

We calculated chlorophyll pathway index values for all three leaf tissues using the normalized transformed expression data of the 22 genes encoding the steps in chlorophyll biosynthesis and the chlorophyll cycle (Figure 1, Figure 4, supplemental Table S5). The Z-score for each gene in each tissue of an individual maize line was calculated as follows:

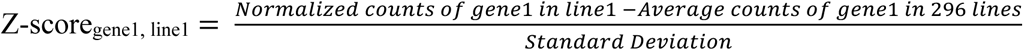

Z-scores of 22 chlorophyll pathway genes were averaged for each tissue type to obtain the chlorophyll biosynthesis index for each inbred line.

### Phenotypic data

Phenotypic data were obtained from previously published studies [25,31]. These included 34 traits from the wild-type and mutant siblings of 343 F_1_ families resulting from crosses between diverse maize lines and *Oy1-N1989/oy1*^B73^:B73. The description of these traits is as follows. Chlorophyll contents were measured at points using a chlorophyll content meter CCM-200 plus (Opti-Sciences, Inc., Hudson, NH). CCMI represents the first time point of chlorophyll measurement at 25-30 days after sowing, and CCMII is the second chlorophyll measurement from plants at 45-50 days after sowing. The isogenic wild-type trait values for every mutant in this F1 population provided a perfect case-control design for this study. We used these two isogenic genotypes to calculate the ratio of mutant to wild type (Ratio CCM) and the difference between wild type and mutant (Diff CCM). Traits measured at maturity included stalk width (SW), plant height or flag leaf height (FlHT), ear height (EaHT), and ear-to-flag-leaf height (Ea2Fl). Days to reproductive maturity were also obtained from previous studies measured as days to 50% anthesis (DTA) and days to 50% silking (DTS) as described previously [31]. Like CCM, we calculated the ratio (mutant/wild type) and difference (wild type - mutant) for the agronomic traits. In addition, the anthesis silking interval (ASI) was calculated as a difference between DTA and DTS.

### Genome-wide association

SNP data for the association panels were obtained from the imputation of the maize HapMap 3.2.1 data [31,56]. The data were filtered to retain only bi-allelic SNPs. The SNPs were recoded so that all the reference alleles were coded as an “A” and the alternate allele as a “T” and then numericalized to speed computation and simplify the interpretation of allelic effects, as described previously [31]. SNPs with a minor allele frequency (MAF) less than 0.05 were removed for each population to reduce false positives. For cis and trans eGWAS and index mapping, data from 296 maize lines were available, and minor allele filtering resulted in 23,913,217 SNPs tested for associations. For the eGWAS analysis, SNPs within 50 kb of each gene’s start or stop codon were classified as cis-acting, and the SNPs located at a distance of 1Mbp or greater were classified as trans-acting. For the phenotypic data on the F1 families, data from 343 F1 families were available, and minor allele filtering for this population resulted in 22,531,621 SNPs tested for association with each trait. Genome-wide associations were performed by modifying the approach taken in switchgrass [57] to adapt the *bigsnpr* package [58] for maize. A preliminary cut-off at a p-value threshold ≤ 1e-4 was selected to minimize excessive false negative rates. GWAS results were viewed using the interactive browser Zbrowse [59]. Annotations for the candidate genes were obtained from several sources, including Gramene, MaizeGDB (https://www.maizegdb.org;[60,61], JGI genome portal (https://genome.jgi.doe.gov/portal/), and Plaza [62]. Arabidopsis homologs were examined using the Thalemine tool in the Araport project [63,64].

### Phylogenetic analysis

The protein sequences of the orthogroup containing the maize *les22* gene (Zm00001d029074) were obtained from the Plaza monocot 4.5 databases [62]. Multiple sequence alignment was performed using clustalW [65] with default parameters (Gap opening penalty: 10 and gap extension cost: 0.20). A maximum likelihood tree was estimated using IQ-TREE version 1.5.5 [66] with substitution model Q.pfam+G4 predicted by Bayesian information criteria (BIC) [67]. A consensus tree was computed using ultrafast bootstrapping with 1000 replicates [68]. The tree was visualized using iTOL v6 [69].

## Acknowledgments

The authors would like to acknowledge the assistance of the staff and leadership at the Agronomy Center for Research and Education, Rosen Center for Advanced Computing, and the Horticulture Greenhouse at Purdue University, West Lafayette, IN. We are grateful to Dilkes lab members for their comments and discussions. We would particularly like to thank everyone who puts extra effort into scientific software documentation and data dissemination that allows research such as this to determine meaning from open-source repositories, including the laboratory of Ed Buckler, USDA ARS. This work was supported by the United States Department of Energy Office of Science (BER) Grants DE-SC0023305 and DE-SC0020368 and National Science Foundation grant IOS-2309932 to B.P.D.. R.S.K was supported by the USDA NIFA Postdoctoral Fellowship award# 2022-67012-36601.

## Author Contributions

A.K, R.S.K, and B.P.D designed the research. A.K and R.S.K performed experiments. A.K, R.S.K and B.P.D analyzed data and wrote the manuscript. A.K, R.S.K, and B.P.D approved the manuscript.

## Supplemental Data

**Figure S1.** Gene Ontology (GO) analysis of differentially expressed genes (DEGs) in *Oy1-N1989/+* mutants. Circle plots show top significant GO terms for biological processes (BP) and cell component (CC) enriched in DEGs in A) *Oy1-N1989/oy1*^B73^:B73 and (B, C) *Oy1-N1989/oy1*^Mo17^:b094 mutants at FDR≤ 0.05. The outer circle depicts the log_2_ fold change of DEGs in *Oy1-N1989/+* mutants as compared to their wild-type (WT) siblings for each enriched GO term. Blue dots indicate genes induced in *Oy1-N1989/+* as compared to WT and gold dots indicate genes repressed in *Oy1-N1989/+*. The color of the inner circle represents the Z-score, and the thickness represents significance of GO term (-log10 (p-value)).

**Figure S2.** Cis-acting regulatory variation at tetrapyrrole biosynthetic pathway genes. A) Distribution of number of genes in tetrapyrrole biosynthetic pathway whose expression was affected by cis polymorphisms in three leaf tissues; B) Distribution of cis-acting SNPs affecting expression of tetrapyrrole biosynthetic genes detected in three leaf tissues.; C) Manhattan plot showing cis-acting SNPs associated with expression of tetrapyrrole biosynthetic pathway genes at p-value <1e-4. The genes that were associated with SNPs at p-value <1e-20 are labelled. The colors indicate cis polymorphisms detected in three leaf tissues: L3Base (Pink), LMAD (Blue) and LMAN (Grey).

**Figure S3.** Cis-acting polymorphisms at genes encoding steps in tetrapyrrole biosynthetic pathway in A) LMAD, B) LMAN and C) L3Base impact chlorophyll content in *Oy1-N1989/+* mutants. Grey dots depict the cis-polymorphism associated with expression of tetrapyrrole pathway genes at p-value < 1e-4. The colored shapes highlight the SNPs that had significant effect on chlorophyll accumulation traits at p-value < 1e-4.

**Figure S4.** SNPs within a 250kb region around *oy1* associated in trans with transcripts of at least two genes encoding steps in tetrapyrrole pathway in LMAD. Light blue and green colors indicate the branches of the tetrapyrrole pathway. The panels on the right depict the direction of effect of *Oy1-N1989* on transcript accumulation in suppressed and enhanced *Oy1-N1989/+* mutants as compared to their respective congenic wild types at unadjusted p-value ≤ 0.05. Dark blue color indicates genes induced in the mutant and gold color indicates genes repressed in the mutant. White indicates that the genes were not differentially expressed at p-value ≤ 0.05.

**Figure S5.** Trans eQTL hotspots associated with the transcripts of eight or more genes encoding steps in tetrapyrrole biosynthesis pathway in LMAD within a 40 kb window. The colors indicate different branches of tetrapyrrole biosynthetic pathway.

**Figure S6.** Trans eQTL hotspots associated with the transcripts of eight or more genes encoding steps in tetrapyrrole biosynthesis pathway in LMAN within a 40 kb window. The colors indicate different branches of tetrapyrrole biosynthetic pathway.

**Figure S7.** Phylogenetic tree for LES22 protein homologs.

**Figure S8.** Transcriptional regulators of pathway level variation in expression of chlorophyll biosynthetic genes. A) Manhattan plot showing the SNPs associated with variation in chlorophyll biosynthesis index (calculated from 22 chlorophyll pathway genes) in three leaf tissues: L3Base (Pink), LMAD (Blue) and LMAN (Grey) at p-value < 1e-4; B) SNPs at *oy1* locus that affect chlorophyll biosynthesis index.

**Figure S9.** Cis-acting polymorphisms at genes encoding steps in tetrapyrrole biosynthetic pathway impact reproductive maturity and growth in *Oy1-N1989/+* mutants. Manhattan plots showing cis-acting SNPs affecting expression of tetrapyrrole biosynthetic genes in LMAD that had significant impact on A) reproductive maturity, B) stalk width, C and D) plant height traits in wild type and *Oy1-N1989/+* F1 plants. Grey dots depict the cis-polymorphism associated with expression of tetrapyrrole pathway genes at p-value < 1e-4 in LMAD and colored shapes highlight the SNPs that had significant effect on phenotypic traits at p-value < 1e-4.

**Figure S10.** Manhattan plots showing cis-acting SNPs affecting expression of tetrapyrrole biosynthetic genes in LMAN that had significant impact on A) reproductive maturity, B) Stalk width, C and D) Plant Height traits in wild type and *Oy1-N1989/+* F1 plants. Grey dots depict the cis-polymorphism associated with expression of tetrapyrrole pathway genes at p-value < 1e-4 in LMAN and colored shapes highlight the SNPs that had significant effect on phenotypic traits at p-value < 1e-4.

**Figure S11.** Manhattan plots showing cis-acting SNPs affecting expression of tetrapyrrole biosynthetic genes in base of third leaf (L3Base) that had significant impact on A) reproductive maturity, B) Stalk width, C and D) Plant Height traits in wild type and *Oy1-N1989/+* F1 plants. Grey dots depict the cis-polymorphism associated with expression of tetrapyrrole pathway genes at p-value < 1e-4 in L3Base and colored shapes highlight the SNPs that had significant effect on phenotypic traits at p-value < 1e-4.

**Table S1.** Differentially expressed genes in (A) *Oy1-N1989/oy1*^B73^:B73 and (B) *Oy1-N1989/oy1*^Mo17^:b094 compared to their isogenic wild-type siblings.

**Table S2.** Complete DEseq2 output for comparison of (A) *Oy1-N1989/oy1*^B73^:B73 and (B) *Oy1-N1989/oy1*^Mo17^:b094 to their respective wild-type siblings.

**Table S3.** Significant GO terms enriched in DEGs in (A) *Oy1-N1989/oy1*^B73^:B73 (B) *Oy1-N1989/oy1*^Mo17^:b094 mutants at FDR≤ 0.05.

**Table S4.** Log2 fold change of genes associated with significant GO terms in (A) *Oy1-N1989/oy1*^B73^:B73, (B) *Oy1-N1989/oy1*^Mo17^:b094 mutants compared to their respective wild-type siblings.

**Table S5.** Log2 fold change of genes encoding steps in tetrapyrrole biosynthesis pathway in *Oy1-N1989/+* mutants relative to their isogenic wild-type siblings.

**Table S6.** Cis polymorphisms associated with variation in expression of genes encoding steps in tetrapyrrole biosynthesis pathway in three leaf tissues.

**Table S7.** Cis SNPs associated with expression variation in tetrapyrrole biosynthesis pathway genes that affect chlorophyll content in mutant and wild-type F1 plants.

**Table S8.** SNPs at the *oy1* locus are linked in trans to the expression of genes encoding steps in the tetrapyrrole biosynthesis pathway.

**Table S9.** Top SNPs associated in trans with expression of genes encoding steps in tetrapyrrole biosynthesis pathway in (A) L3Base, (B) LMAD, and (C) LMAN.

**Table S10.** A) Trans eQTL hotspots affecting the expression of eight or more genes encoding steps in tetrapyrrole biosynthesis pathway within a 40 kb window in (A) L3Base, (B) LMAD, and (C) LMAN.

**Table S11.** Proportion of trans polymorphisms associated with expression of each gene in tetrapyrrole biosynthesis pathway present in trans-eQTL hotspots in three leaf tissues.

**Table S12.** SNPs present in trans-eQTL hotspots that affect chlorophyll content in mutant and wild-type F1 plants.

**Table S13.** SNPs associated with variation in Chlorophyll Biosynthesis Index at p<1e-4 in three leaf tissues.

**Table S14.** Cis SNPs associated with expression variation in tetrapyrrole biosynthesis pathway genes that affect plant height traits in mutant and wild-type F1 plants at p-value <1×10^-4^.

**Table S15.** Effect of *Oy1-N1989/+* on transcript levels of genes encoding steps in (A) Strigolactone biosynthesis, (B) Gibberellin biosynthesis and signaling, (C) Brassinosteroid biosynthesis and signaling, and (D) Auxin biosynthesis and signaling.

